# Multiple *in vivo* roles for the C-terminal domain of the RNA chaperone Hfq

**DOI:** 10.1101/2021.08.13.456236

**Authors:** Kumari Kavita, Aixia Zhang, Chin-Hsien Tai, Nadim Majdalani, Gisela Storz, Susan Gottesman

## Abstract

Hfq, a bacterial RNA chaperone, stabilizes small regulatory RNAs (sRNAs) and facilitates sRNA base-pairing with target mRNAs. Hfq has a conserved N-terminal domain and a poorly conserved disordered C-terminal domain (CTD). In a transcriptome-wide examination of the effects of a chromosomal CTD deletion (Hfq_1-65_), the *Escherichia coli* mutant was most defective for the accumulation of sRNAs that bind the proximal and distal faces of Hfq (Class II sRNAs), but other sRNAs also were affected. There were only modest effects on the levels of mRNAs, suggesting little disruption of sRNA-dependent regulation. However, cells expressing Hfq lacking the CTD deletion in combination with a weak distal face mutation were defective for the function of the Class II sRNA ChiX and repression of *mutS*, both dependent upon distal face RNA binding. Loss of the region between amino acids 66-72 was critical for this defect. The CTD region beyond amino acid 72 was not necessary for distal face-dependent regulation, but was needed for functions associated with the Hfq rim, seen most clearly in combination with a rim mutant. Our results suggest that the C-terminus collaborates in various ways with different binding faces of Hfq, leading to distinct outcomes for individual sRNAs.

## INTRODUCTION

Bacteria use small regulatory RNAs (sRNAs) as part of the response to environmental stresses, repressing outer membrane protein synthesis, remodeling metabolism, and affecting synthesis of important transcription factors and virulence factors [reviewed in (1–3)]. Many of these sRNAs regulate translation and decay of their target mRNAs through limited-complementary base pairing with the aid of the RNA chaperone Hfq [reviewed in (4)].

Hfq was first identified in *E. coli* as a <u>h</u>ost factor required for bacteriophage <u>Q</u>β RNA replication (5). This homo-hexameric protein is a member of the family of Sm and Sm-like RNA binding proteins (6). Hfq uses multiple RNA binding faces to perform flexible matchmaking between sRNAs and mRNAs [reviewed in (7)]. The proximal face of Hfq has specificity for U-rich regions and is essential for binding of all Hfq-dependent sRNAs (8). The distal face can recognize and bind repeated AAN sequences (8, 9). The rim site has acidic patches of positively-charged arginines, which are ideal for binding negatively-charged RNA [reviewed in (10)] and have been found to interact with A/U rich sequences (11). The 102- amino acid *E. coli* Hfq has an evolutionarily conserved N-terminal domain from residues 1 to 65. The flexible and less conserved C-terminal domain (CTD, residues 66-102) is not present in most published crystal structures and has been predicted to be unstructured (12). In a co-crystal structure of full length Hfq with the RydC sRNA, residues beyond 70 were poorly ordered, but appeared to be making distributive contacts over the sRNA surface (13).

While the traditional view is that only proteins with well-defined structure have function, intrinsically disordered regions recently have been shown to have crucial biological roles [reviewed in (14)]. In some cases, binding to a partner or ligand is coupled to folding; in other cases, the disordered region can act as a flexible linker for macromolecule assembly [reviewed in (14)]. Therefore, it seemed possible that the Hfq CTD has important functions.

The CTD of Hfq has been studied in the past, but some studies reported that it is dispensable for Hfq function while others found it is critical for sRNA action [see, for instance, (15–18)]. An argument for a function is that many bacteria have maintained long Hfq C-terminal tails during evolution, suggesting that the CTD confers benefits to these bacteria (4). One study showed that the CTD plays a role in the release of sRNA-mRNA duplexes and drives sRNA competition on Hfq *in vitro* and *in vivo* (18). Our current model of Hfq function for sRNA regulation is that the Hfq ring promotes annealing of a bound sRNA and complementary bound mRNA, and then releases the paired duplex, thus regulating the fate (translation and stability) of the mRNA (8, 19). The sRNA and mRNA must bind to different surfaces of Hfq for this to occur. We found that sRNAs could be classified based on which faces of Hfq they bind. Class I sRNAs bind to proximal and rim faces, and pair with targets that bind the distal face, while Class II sRNAs bind to proximal and distal faces, and pair with targets that bind to the rim (8). Most sRNAs are stabilized by binding to Hfq. After pairing, however, Class I sRNAs and some Class II sRNAs are generally released from Hfq along with their targets, and are rapidly degraded. If the mRNA targets for a given sRNA cannot bind (due to mutation in the Hfq binding site for those targets), the sRNA is stabilized and will accumulate to higher levels in the cell (8). A subset of Class II sRNAs are not released after pairing but are believed to be used for multiple rounds of pairing and mRNA degradation (8, 20). Disruption of stable Hfq binding can decrease their stability and therefore lead to reduced accumulation of these Class II sRNAs. Thus, levels of sRNAs are affected in multiple ways by their interactions with Hfq, and Hfq mutants can have profound effects on both sRNA function and sRNA levels, though these are not always directly correlated.

Here we first examined the global effect of the Hfq CTD on RNA levels. We then used combinations of mutations in the Hfq CTD and mutations in the RNA binding faces of Hfq to examine the ways in which the CTD interacts with sRNAs to affect their accumulation and function. We find that a deletion of the CTD does not result in dramatic phenotypes in *E. coli.* However, combinations of deletions or mutations in the CTD with mutations in the RNA binding faces of Hfq show strong synergistic effects, possibly suggesting that the CTD collaborates with these surfaces. Interestingly, different parts of the CTD play distinct roles for individual sRNAs, reflected in changes in the accumulation and functions of these sRNAs, providing an explanation for previous discrepant results.

## MATERIALS AND METHODS

### Bacterial strains, plasmids, and strain construction

The bacterial strains used in this study are listed in Supplementary Table S1; they are derivatives of *Escherichia coli* K12 strain MG1655 (Fred Blattner collection). Oligonucleotides and gblocks used for strain construction are in Supplementary Table S2. Plasmids used in this study are: vector control (pBR-plac), pBR-ChiX, pBR-RyhB (8).

Hfq alleles were first constructed in the chromosome in strains DJS2255 or DJS2814 (Table S1) by λ-red recombineering (21), and confirmed by sequencing. The recipient strains contain a Kan^R^-pBAD-*ccdB* cassette in place of the *hfq* gene and miniλ::*tet*, encoding lambda recombination functions repressed by the λ*cI*857 repressor. CcdB is toxic, and thus was used as a counter-selectable marker during recombineering. Strains were first grown in 10 ml LB with 1% glucose to an OD_600_∼ 0.6-0.8 at 32°C to keep the level of expression of CcdB very low. Expression of the recombinase genes from miniλ was induced by shifting the culture flask to 42°C for 15 min, after which cells were rapidly cooled by shaking the flask in an ice slurry for 2 min, followed by a further 10 min on ice. Cells were collected by centrifugation at 4,500 rpm at 4°C for 10 min. The cell pellet was washed three times with an equal volume of ice-cold sterile water and then resuspended in 100 μl of ice-cold sterile water. 50 ng of DNA was added to 50 µl of cells using electroporation. Cells were allowed to recover in LB+1% glucose for 1 h. Cells were kept at room temperature overnight to improve the recombination frequency. 100 µl of cells were plated on LB with 1% arabinose and incubated overnight at 37°C. After restreaking for purification on LB with 1% arabinose and then on LB, loss of the kanamycin marker was confirmed, and the final strain confirmed by sequencing of *hfq*. The alleles were then transferred by P1 transduction to reporter strains used for assays, using a recipient with the appropriate reporter and Δ*purA::kan* Δ*hfq::cat-sacB*, selecting PurA^+^ colonies and screening for loss of the chloramphenicol resistance marker in the linked *hfq* gene (see Table S1).

### Bacterial growth conditions

All strains used in this study were grown at 37°C under aerobic conditions in LB (KD Medical, MD). Where required, the liquid and solid media were supplemented with ampicillin (100 μg/ml).

The bacteria used for the RNA-Seq experiment, WT (KK2440) and Hfq_1-65_ (KK2448), were grown to OD_600_ ∼ 1.0 in LB.

### RNA co-immunoprecipitation with Hfq and isolation of total RNA

Polyclonal antibodies to Hfq were generated previously by immunizing rabbits with purified Hfq protein (Covance). RNAs that co-immunoprecipitated (co-IP) with Hfq were isolated as described previously (22) with the following modifications. Overnight cultures of WT (KK2440) and Hfq_1-65_ (KK2448) cells were grown to OD_600_ ∼ 1.0 in LB medium. Cells corresponding to the equivalent of 40 OD600 were collected, and lysates were prepared by vortexing cells with 212-300 μm glass beads (Sigma) in a final volume of 2 ml of lysis buffer (20 mM Tris-HCl/pH 8.0, 150 mM KCl, 1 mM MgCl_2_, 1 mM DTT). IPs were carried out using 200 μl of anti-Hfq antibody, 240 mg of protein A-Sepharose CL-4B (GE Healthcare), and 1.9 ml of cell lysate. Co-IP RNA was isolated from protein A-Sepharose beads by extraction with phenol: chloroform:isoamyl alcohol (25:24:1), followed by ethanol precipitation. Total RNA was isolated from 100 μl of cell lysate by Trizol (Thermo Fisher Scientific) extraction followed by chloroform extraction and isopropanol precipitation. Total and co-IP RNA samples were resuspended in 30 μl and 50 μl of DEPC H_2_O, respectively.

Note that we previously found that the anti-Hfq antibody used for immunoprecipitation recognizes the truncated Hfq_1-65_ significantly less well than the full length Hfq (18). We did not normalize immunoprecipitation efficiencies. However, we kept this difference in mind when interpreting any comparisons between the wild-type and *hfq*_1-65_ immunoprecipitation samples. Presumably this difference would lead to overestimation of the IP defect of *hfq*_1-65_ versus *hfq*^+^, leading us to focus somewhat more on the total RNA levels in this study.

### RNA-Seq library construction

RNA samples (400 ng) were used for cDNA library preparation. Library construction was carried out based on the RNAtag-Seq methodology (23), which was adapted to capture bacterial sRNAs (24). The index primers used in this study are listed in Supplementary Table S2. To enrich the libraries, 12 cycles of PCR were carried out at the end step of enrichment. Bead purification was performed twice at the last step to ensure high purity of the library from the adaptor dimer. The sequencing was done using an Illumina HiSeq2500 platform.

### RNA-Seq data analysis

The Illumina reads in FASTQ format were first trimmed by Trimmomtaic (25) version 0.36 to remove the adaptor sequences and low quality reads. The RNA-Seq pipeline READemption (26) version 0.4.5. was then used to process those trimmed reads. Reads longer than 12 nucleotides were aligned to the *Escherichia coli* str. K-12 substr. MG1655 reference sequence (NC_000913.3) from the NCBI by using segemehl (27) version 0.2.0. The “deseq” subcommand in READemption executing DESeq2 version 1.20.0 was used, along with a fold-change of 2.0 or higher and an adjusted P value below 0.1 as the thresholds to find differentially expressed genes. The data are summarized in Tables S3, S4, and S5.

### Western blot analysis

Protein samples were generated by collecting the pellet for 1 ml of cells at OD_600_ ∼ 1.0 and lysing with 100 µl 1X SDS sample buffer (New England Biolabs) and DTT by boiling for 10 min. The protocol is based on that described previously (28). Samples (10 µl) were resolved on a 12% bis-Tris polyacrylamide NuPAGE gel (Invitrogen, USA) in 1X of 2-(N-morpholino)ethanesulfonic acid (MES) buffer (Invitrogen, USA) for 35 min at 180V. The gel was transferred using the iBlot2 (ThermoFisher Scientific) to a nitrocellulose membrane, blocked with 3% milk in 1X PBST (PBS with 0.1% Tween 20) and treated with primary anti-Hfq antibody, diluted 1:5000 in 0.3% milk in 1x PBST; the anti-Hfq antibody was preadsorbed with cell extract from a strain deleted for *hfq*. After washing, the membrane was incubated with 1:1000 dilution of goat anti-rabbit AP-linked secondary antibody (Cell Signaling) in 0.3% milk in 1XPBST at room temperature for 1hr, washed and developed using CDPStar (Invitrogen).

### β-galactosidase assays (Miller assay)

Cells expressing arabinose-inducible (P_BAD_) or constitutively-transcribed (Cp12b) translational fusions were grown in LB at 37°C, at 250 rpm; samples were collected and assayed for β- galactosidase activity using o-nitrophenyl-β-D-galactopyranoside (ONPG) as substrate as per the standard method (29). For the P_BAD_*-chiP*-*lacZ* translational fusion, 0.002% arabinose was added to LB, cells were grown to OD_600_ ∼ 1.0, and assayed. For Cp12b*-mutS-lacZ*, cells were grown until late stationary phase in LB and assayed (30). For P_BAD_*-sodB-lacZ* and P_BAD_*-chiPsodB-lacZ* strains carrying the pBR-RyhB plasmid, cells were grown to early stationary phase in LB with 100 μg/ml ampicillin, 0.002% arabinose, and 10 μM IPTG as previously described; Δ*fur* strains containing P_BAD_*-sodB-lacZ* were treated similarly, except that no plasmid was present and thus ampicillin was not included (8). All assays were performed at least in triplicate, and the mean and standard deviations are presented in each figure. Unless indicated otherwise in the figure legend, three independent cultures were assayed.

### Northern blot analysis

Equal amounts (5 μg) of total RNA were fractionated on 8% polyacrylamide urea gels containing 6 M urea (1:4 mix of Ureagel Complete to Ureagel-8 (National Diagnostics) with 0.08% ammonium persulfate) in 1X TBE buffer at 300V for 90 min. The RNA was transferred to a Zeta-Probe GT membrane (Bio-Rad) at 20 V for 16 h in 0.5X TBE and crosslinked to the membranes by UV irradiation. RiboRuler Low Range RNA ladders (Thermo Fisher Scientific) run on gel were identified by UV-shadowing. The membranes were then blocked in ULTRAhyb-Oligo Hybridization Buffer (Ambion) for 2 h at 45°C. Oligonucleotide probes (listed in Supplementary Table S2) were 5′ ^32^P-end labeled with 0.3 mCi of γ-^32^P ATP (Perkin Elmer) by incubating with 10 U of T4 polynucleotide kinase (New England Biolabs) at 37°C for 1 h. Subsequently, labeled probes were purified using Illustra MicroSpin G-50 columns (GE Healthcare) and 40 pmol of these probes was added to the blocked membranes. After an overnight incubation, the membranes were rinsed twice with 2X SSC/0.1% SDS at room temperature, once with 0.2X SSC/0.1% SDS at room temperature, washed for 25 min with 0.2X SSC/0.1% SDS at 45°C, followed by a final rinse with 0.2X SSC/0.1% SDS at room temperature prior to exposure to X-ray film. For sequential probing, blots were stripped by three 7 min incubations in boiling 0.2% SDS followed by three 7 min incubations in boiling water.

## RESULTS

### Global examination of the role of the Hfq CTD

In previous work, we had examined the effect of deleting the Hfq CTD on specific sRNAs (Class I sRNAs RyhB, GcvB, OmrB, ArcZ and Spot 42 and Class II sRNAs ChiX, MgrR, CyaR and McaS) by expressing the truncated protein Hfq_1-65_ from the chromosomal locus. We found that loss of the CTD resulted in decreased stability and lower levels of the Class II sRNAs, which bind the proximal and distal faces of Hfq, while only OmrB was strongly affected among the Class I sRNAs tested (18). Here we obtained a global view of the role of the CTD by comparing WT and Hfq_1-65_ strains by RNA-Seq both for the whole transcriptome and for RNA collected after Hfq immunoprecipitation (IP). Cells were grown in LB and samples were collected at OD_600_ ∼ 1; the experiment was conducted with two biological replicates and the data were analyzed by READemption and DESeq2. There was good reproducibility between the replicates (Supplementary Figure S1).

The volcano plot of specific RNA levels in Hfq_1-65_ vs WT Hfq is shown in Figure 1A, and the volcano plot for the co-IP experiment is shown in Supplementary Figure S2A. Supplementary Table S3 contains the RNA-Seq and co-IP results for all genes in the WT and Hfq_1-65_ strains. Genes that showed more than 2-fold increased or decreased total RNA levels in the Hfq_1-65_ background compared to WT are listed in Supplementary Table S4 and those with changes of 2-fold or more upon co-IP with Hfq_1-65_ compared to WT are listed in Supplementary Table S5. We also examined the enrichment of each gene by co-IP compared to total RNA in the WT background (Supplementary Tables S3, S4 and S5, column V). Here we focus our discussion on those genes that are enriched for Hfq binding (> 2-fold), since they are more likely to be directly affected by the changes in Hfq; these are highlighted in yellow in Tables S4 and S5, columns A and V.

**Figure 1.**
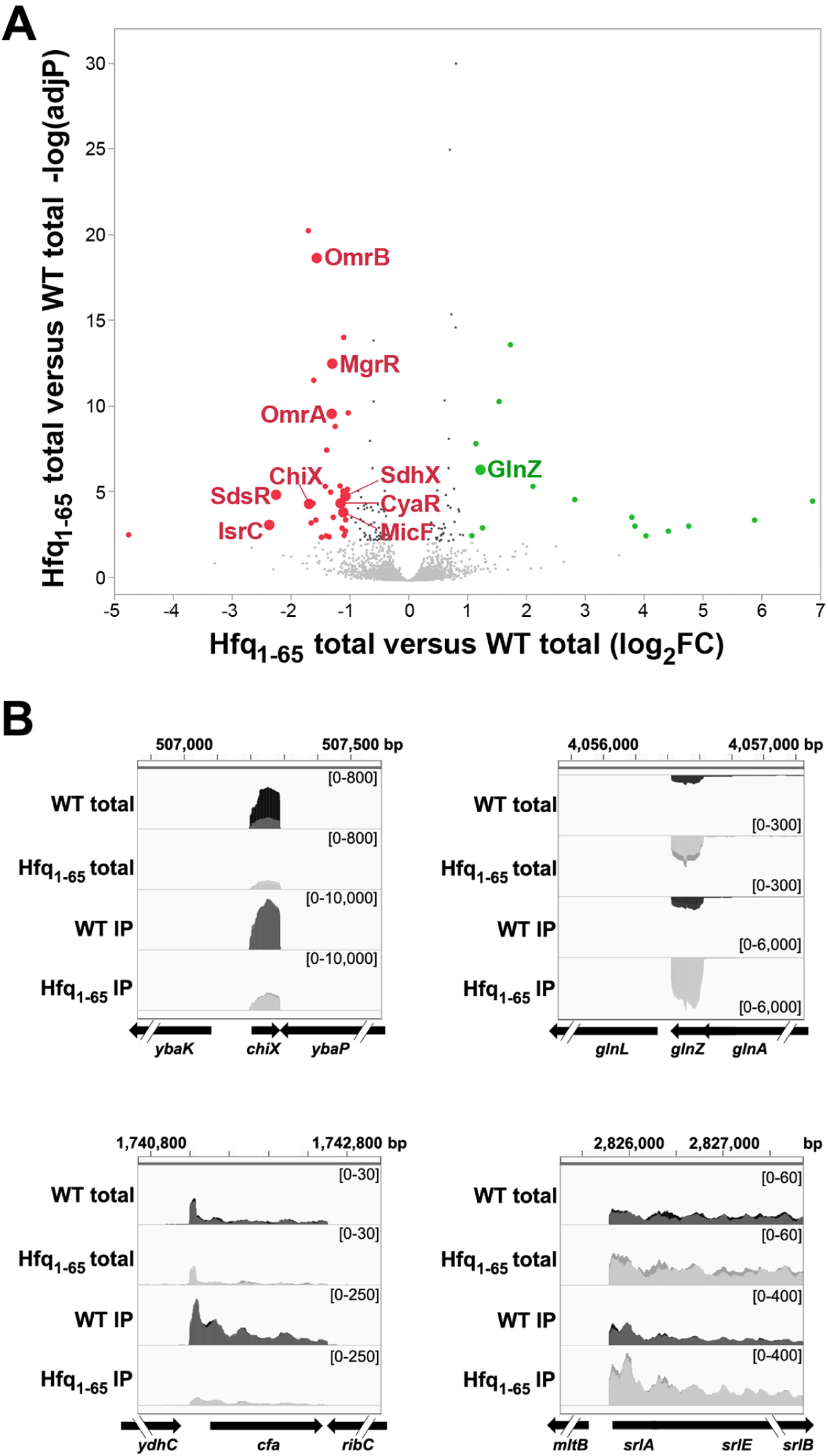
Effect of CTD deletion on sRNA and mRNA levels. (**A**) Statistical analysis and visualization of RNAs levels for total RNA isolated from Hfq WT and Hfq_1-65_. Volcano plot comparing total RNA from Hfq_1-65_ (KK2448) to Hfq WT(KK2440). Red dots are those that were downregulated more than 2-fold in Hfq_1-65_, while green dots were upregulated by more than 2- fold. Some sRNAs are labeled. All genes with adjusted P values greater than 0.1 are in grey (4296 genes); genes with adjusted P values < 0.1, and fold change of less than <2x change (100 genes) are shown in black. (**B**) Integrative Genomics Viewer [IGV_27.2; (47)] images of two sRNAs (ChiX and GlnZ) and two mRNAs (*cfa* and *srl* operon) that show significant changes in levels in a comparison of total and IP RNA for Hfq_1-65_ (KK2448) to Hfq WT(KK2440). GlnZ is the only sRNA that accumulates to higher levels in the absence of the Hfq CTD.

### Lack of Hfq CTD also affects some Class I sRNAs

In Table S4B, 13 Hfq-binding RNAs showed >2-fold decreases in abundance in Hfq_1-65_ compared to WT Hfq; eight of these are known Hfq-binding sRNAs (labeled sRNA in column B, Type). Three of the five others are enriched in WT IP in a pattern that suggests they too may be sRNAs encoded at the 3′ UTRs of mRNA genes (labeled sRNA* in column B, see below). In Table S5, enrichment of the RNAs by IP with Hfq increased our ability to detect less well-expressed sRNAs. 51 RNAs showed both a >2-fold decrease in Hfq_1-65_ compared to WT Hfq (listed in Table S5B), and 2-fold or better enrichment in binding to Hfq. 17 of these are known Hfq-binding sRNAs (Table S5B). These include both Class I (OmrB, DsrA, OmrA, MicF, GadY, DicF, Spot 42, MicA) and Class II (CyaR, MgrR, ChiX) sRNAs. Some signals with >2- fold decreases also correspond to recently defined sRNAs encoded at the 5′ and 3′ UTR of genes such as MotR (corresponding to the 5′ of *motA*) and UhpU (corresponding to the 3′ of *uhpT*) (31), as well as others such as *tdcG* and *ygaM* that have RNA-Seq signals suggesting they too likely are 3′ UTR-derived sRNAs (indicated as sRNA*). The RNA-Seq browser image for ChiX, previously shown to be decreased in Hfq_1-65_, is shown in Figure 1B; those for MotR, UhpU as well as the 3′ UTRs of *tdcG* and *ygaM* are shown in Figure S2B.

Only one Hfq-binding sRNA, the 3′ UTR-derived sRNA GlnZ (32, 33) showed 2-fold higher levels in both total RNA and co-IP with Hfq_1-65_ compared to WT Hfq (Figure 1B). The results generally support and extend our previous findings (18), but demonstrate that the Hfq CTD is likely to stabilize at least some Class I sRNAs in addition to the previously documented effects on the Class II sRNAs.

### mRNAs affected by the Hfq CTD generally are targets for affected sRNAs

The Hfq-binding transcripts annotated as encoding proteins and that showed log_2_-fold lower co-IP with Hfq_1-65_ compared to WT Hfq (Supplementary Table S5B) include some genes known to be subject to sRNA-dependent regulation (34–36). For example, *cfa* (Figure 1B) is positively regulated by RydC and ArrS (36). For this mRNA, a decrease in levels of the sRNAs that positively regulate it may be the major determinant for the decreased levels. Other mRNAs in this group with lower co-IP with Hfq_1-65_ compared to WT Hfq may similarly be subject to positive regulation by Hfq-binding sRNAs.

Only two Hfq-binding mRNAs showed >2-fold higher co-IP with Hfq_1-65_ compared to WT Hfq (Supplementary Table S5A). Both are associated with strong negative regulation by sRNAs. *chiP* is strongly repressed by ChiX and that repression is somewhat relieved in the Hfq_1- 65_ strain [(18), Supplementary Table S5A, see below]. The other mRNA, *srlA* (Figure 1B), which is reasonably abundant under the growth conditions used here (Supplementary Table S4), is a known target for repression by Spot 42 (37). Thus, here too, the changes in mRNA signals are likely secondary to changes in the levels of the sRNAs. Overall, it is noteworthy that only a very small subset of the many mRNA targets subject to sRNA regulation are sufficiently perturbed to result in a 2-fold or better change in either the total RNA or the IP signals in cells expressing Hfq_1-65_. While parallel RNA-Seq experiments have not been done with other *hfq* mutants, older microarray studies show dozens of genes increased 2-fold or better in cells carrying Q8A, an allele that reduces or abolishes sRNA binding to Hfq and thus may abrogate the effects of many sRNAs on mRNAs (38). Thus, the lack of an effect observed for strains expressing Hfq_1-65_ suggests that the loss of the CTD does not strongly affect Hfq function *in vivo*.

### Functional analysis of the Hfq CTD in cooperation with binding face mutations

The global analysis above suggests that the loss of the CTD perturbs the levels of some Hfq-dependent sRNAs, resulting in very modest effects on mRNA targets. If the role of the CTD is somewhat redundant with roles of other parts of Hfq, it seemed possible that combining the CTD deletion with the previously studied mutations in binding faces of Hfq would allow us to further define *in vivo* roles for the CTD. We thus created multiple isogenic strains, mutant for one or another of the Hfq faces in combination with the CTD deletion (Figure 2A). By immunoblot analysis, the levels of the Hfq derivatives with the CTD deletion appear lower than the full-length derivatives (Supplementary Figure S3). This is likely due, in part, to epitopes present in the flexible CTD recognized by the polyclonal antiserum and thus missing in the deletion derivative (18). Some Hfq mutants, particularly Hfq_1-65_R16A and Hfq_1-65_Y25D, may be unstable, although Hfq_1-65_R16A is clearly functional in some assays (see below).

**Figure 2.**
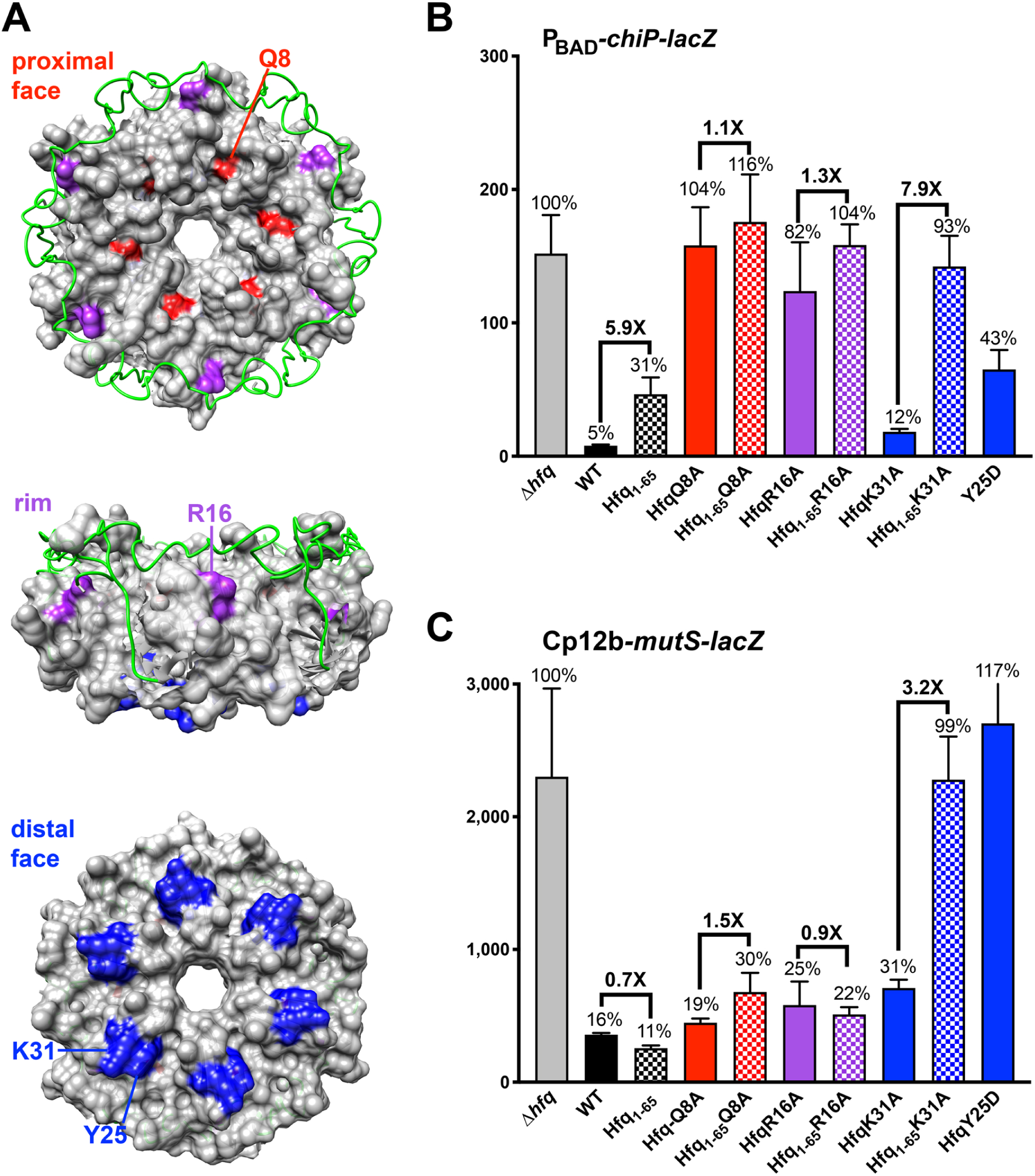
Synergy between Hfq CTD and weak distal face mutant (K31A) of Hfq is critical for distal face regulation. (**A**) Hfq structure (PDB:4V2S) showing mutations on the different faces (proximal Q8A in red, rim R16A in purple, distal Y25D or K31A in blue, modeled CTD in green). Structural images of Hfq were produced using the UCSF Chimera (48) package from the Resource for Biocomputing, Visualization, and Informatics at the University of California, San Francisco. (**B**) β-galactosidase activity measured in P_BAD_-*chiP-lacZ* strains, where ChiX is expressed from the chromosome. Strains used (Supplementary Table S1) are isogenic derivatives all carrying *lacI’::P_BAD_-chiP-lacZ* and *hfq* alleles at the native locus [Δ*hfq* (DJS2689), WT (DJS2690), *hfq_1-65_* (KK01), *hfqQ8A* (DJS2691), *hfq_1-65_Q8A* (KK2414), *hfqR16A* (DJS2693), *hfq_1-65_R16A* (KK02), *hfqK31A* (DJS2695) and *hfq_1-65_K31A* (KK04) and *hfqY25D* (DJS2694)]. Strains were grown in LB media with 0.002% arabinose at 37°C to early stationary phase (OD_600_ ∼ 1.0) and assayed for β-galactosidase activity as described in Materials and Methods. Data are average of three independent assays and error bars represent standard deviation of mean. The *hfqQ8A* (DJS2691) and *hfq_1-65_Q8A* (KK2414) samples were assayed in a separate experiment from the other mutants. (**C)** Effect of the Hfq CTD on regulation of *mutS*. β-Galactosidase activity measured for a *mutS*-*lacZ* fusion, which is known to be directly regulated by Hfq in the absence of sRNAs. The fusion is expressed from the constitutive Cp12b promoter. Isogenic strains all contain *lacI’::kan-Cp12b-mutS-lacZ* and *hfq* mutations at the native locus [Δ*hfq* (KK2455), WT (KK2456), *hfq_1-65_* (KK2457), *hfqQ8A* (KK2460), *hfq_1-65_Q8A* (KK2461), *hfqR16A* (KK2462), *hfq_1-65_R16A* (KK2463), *hfqK31A* (KK2466), *hfq_1-65_K31A* (KK2467) and *hfqY25D* (KK2464)]. Strains were grown at LB at 37°C to late stationary phase (OD_600_ ∼ 3) and assayed as above. Data are the average of three assays, two from one culture and one from an independent culture, and error bars represent the standard deviation of the mean.

### Synergistic effect of CTD deletion and distal face mutations for Class II sRNA-dependent regulation

The Class II sRNA ChiX is a strong negative regulator of *chiP* expression (39). ChiX is expressed from its native gene even in the absence of any specific inducing conditions, and thus expression from chromosomal *chiX* is sufficient for effective Hfq-dependent repression of *chiP* in wild-type (WT) cells (Figure 2B, compare WT to Δ*hfq*). We previously found that ChiX, normally very stable, becomes somewhat unstable in Hfq_1-65_ cells and accumulates to about half the level found in WT cells (18); this difference in accumulation was also observed in our RNA-Seq data (Figure 1A and B; Table S4). A *chiP*-*lacZ* translational fusion also was regulated somewhat less effectively by the Hfq_1-65_ mutant. Single mutations in the faces of Hfq were tested, either alone or in combination with the Hfq_1-65_ truncation, for ChiX repression of the *chiP-lacZ* fusion. A mutation in the proximal face of Hfq (Q8A) interfered significantly with ChiX binding and stability (38), and thus abrogated regulation completely (Figure 2B). The rim mutation R16A also was defective, in this case likely because the mRNA target binds to the rim (8). Combination of either of these mutations with the CTD deletion resulted in similar defects (Figure 2B).

A weak distal face mutant (K31A) had little or no effect on ChiX repression, consistent with previous results (38), but combining K31A with the Hfq_1-65_ deletion led to a complete loss of function for regulation of *chiP* (Figure 2B, compare blue bar to checked blue bar). Hfq protein was not detected for the combination of a stronger distal face mutant, Y25D, with Hfq_1-65_ (Supplementary Figure S3), and thus this double mutant was not further studied. If the lack of regulation by Hfq_1-65_K31A is due to the instability of ChiX, as seen previously for Hfq_1-65_ (18), overproducing the sRNA might bypass the regulatory defect. This was tested qualitatively in a plate assay (Supplementary Figure S4). Multicopy ChiX did not suppress the regulatory defect in a strain deleted for *hfq* (black arrow) or the mutant in mRNA binding, R16A, either by itself or in combination with Hfq_1-65_. However, overproduction of ChiX was sufficient to repress *chiP* in the Hfq_1-65_K31A double mutant (red arrow, right-hand plate, compared to left-hand plate, expressing vector). The suppression by multicopy ChiX also suggests that the Hfq_1-65_K31A protein is at least partially functional.

### Synergistic effect of CTD deletion and distal face mutations for sRNA-independent regulation

In the assay of ChiX regulation of *chiP* (Figure 2B), we suggest that the defect in regulation with the Hfq_1-65_K31A double mutant is likely due to less stable binding of ChiX to the distal face of Hfq. If loss of the Hfq CTD acts synergistically with weak distal face mutants, we might expect to see a similar synergistic effect for mRNA binding to the distal face. The Hfq distal face has been reported to repress *mutS* mRNA directly without the involvement of sRNAs; mutations in the sRNA binding face (Q8A) did not block this regulation, but a strong mutation in the distal face (Y25D) relieved repression (30) (Figure 2C). We took advantage of this system to evaluate the role of the Hfq CTD for sRNA-independent regulation by testing the Hfq_1-65_ single and double mutants for negative regulation of a *mutS-lacZ* fusion. For this assay, Hfq_1-65_ itself was as effective as WT. While the Y25D mutant was fully defective, K31A had only a modest effect on the ability of Hfq to repress the *mutS* fusion. However, the Hfq_1-65_K31A double mutant was fully defective for *mutS* repression, as it was for repression of *chiP* (Figure 2B and 2C). These results suggest a significant role of the Hfq CTD in reinforcing or protecting binding to the distal face of Hfq. This is consistent with and extends our previous results on the effect of the Hfq CTD on Class II sRNAs, all of which (by definition), bind to the distal face (18). The effective repression of *mutS* in the Hfq_1-65_Q8A and Hfq_1-65_R16A double mutants also indicates that these double mutants, even if present at lower levels (Figure S3), are functional for some activities (compare repression to loss of regulation in strain deleted for *hfq*, Figure 2C).

### ChiX-mediated regulation requires arginine 66 of Hfq CTD

*E. coli* Hfq is 102 amino acids long, and the truncation used in the experiments above and in a variety of previous studies (16,18,40) shortens it to 65 amino acids. We were interested in defining what part of the CTD is critical for its functions related to the distal face. Thus, derivatives of Hfq expressing truncations of 10, 20, and 30 amino acids (Hfq_1-92_, Hfq_1-82_, and Hfq_1-72_) were created by replacing the WT *hfq* gene in the chromosome. *In vitro* studies by Santiago-Frangos et al. implicated the acidic tip of Hfq (amino acids 98-102, EETE), and the region just upstream of that (amino acids 88-97, called by them the linker) in Hfq function (17). Derivatives of Hfq in which these regions were mutated were also created (Figure 3A).

**Figure 3.**
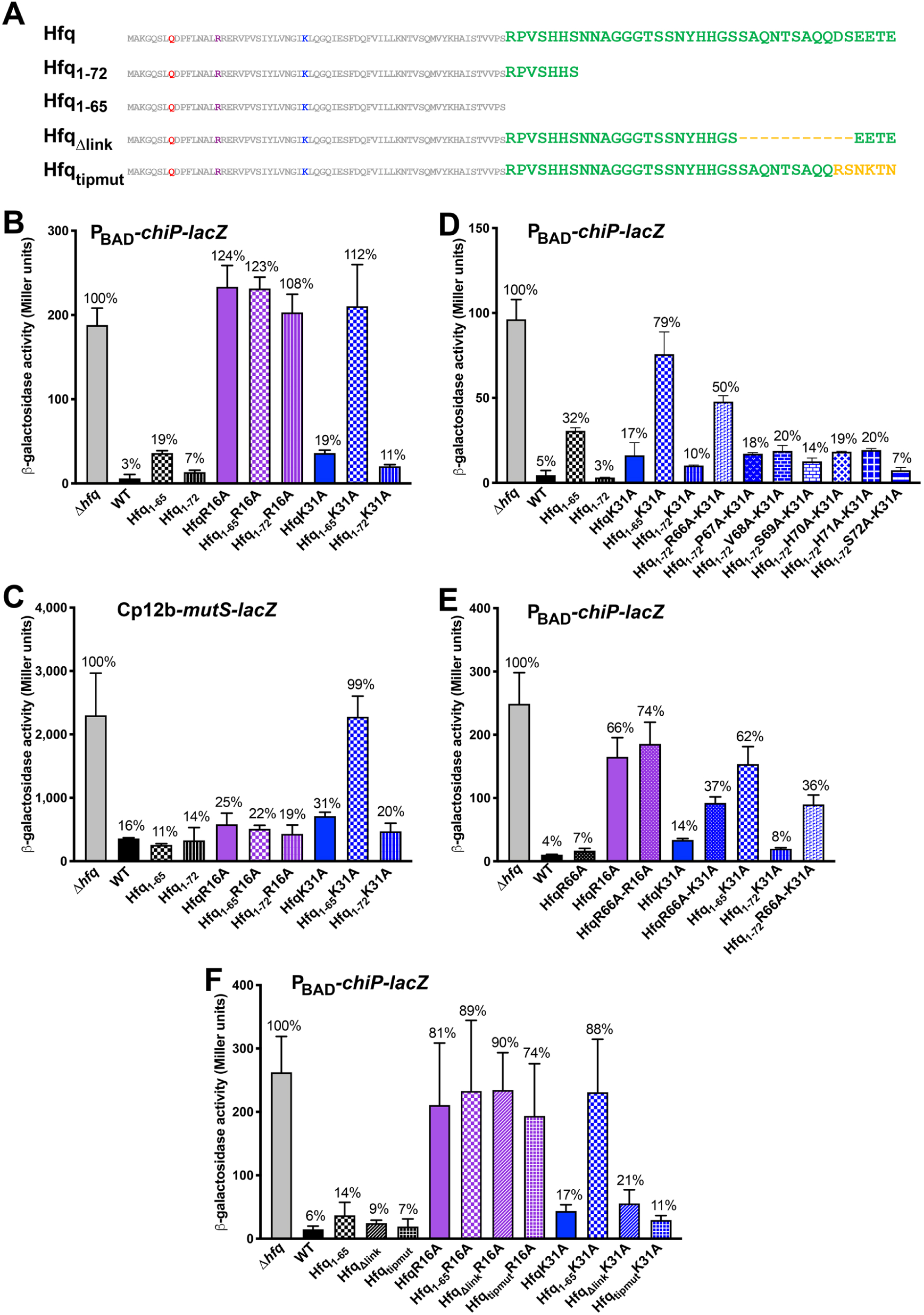
Dissecting the Hfq CTD for distal-face regulation. (**A**) Schematic of Hfq CTD truncations and mutations. **(B)** β-galactosidase activity measured in isogenic strains carrying the P_BAD_-*chiP-lacZ* fusion, CTD truncations Hfq_1-65_ and Hfq_1-72_ with mutations in rim and distal face of Hfq [Δ*hfq* (DJS2689), WT (DJS2690), *hfq_1-65_* (KK01), *hfq_1-72_* (KK2438), *hfqR16A* (DJS2693), *hfq_1-65_R16A* (KK02), *hfq_1-72_R16A* (KK2657), *hfqK31A* (DJS2695), *hfq_1-65_K31A* (KK04) and *hfq_1-72_K31A* (KK2658)]. Strains were grown and assayed for β-galactosidase activity as in Figure 2B. Data are the average of four assays, two from each of two independent cultures (except for a single biological sample for *hfq*_1-72_), and error bars represent the standard deviation of the mean. (**C**) β-galactosidase activity measured in Cp12b-*mutS-lacZ* directly regulated by Hfq, CTD truncations Hfq_1-65_ and Hfq_1-72_ with mutations in rim and distal face of Hfq [Δ*hfq* (KK2455), WT (KK2456), *hfq_1-65_* (KK2457), *hfq_1-72_* (KK2689), *hfq-R16A* (KK2462), *hfq_1-65_R16A* (KK2463), *hfq_1-72_R16A*(KK2691), *hfqK31A* (KK2466), *hfq_1-65_K31A* (KK2467) and *hfq_1-72_K31A* (KK2692)] expressed from the chromosome. The three strains with Hfq_1-72_ alone or in combination were assayed as a group as in Figure 2C, and values for all but the Hfq_1-72_ derivatives were from the data in Figure 2C. Data are the average of three assays, two from one culture and one from an independent culture, and error bars represent the standard deviation of the mean. (**D**) β-galactosidase activity measured in isogenic strains carrying the *lacI’::P_BAD_-chiP-lacZ* fusion, with single alanine substitutions for positions 66-72 of the CTD, combined with K31A in Hfq_1-72_, were expressed from the native *hfq* site [Δ*hfq* (DJS2689), WT (DJS2690), *hfq_1-65_* (KK01), *hfq_1-72_* (KK2438), *hfq_1-65_K31A* (KK04), *hfq_1-72_K31A* (KK2658), *hfq_1-72_R66A-K31A* (KK2680), *hfq_1- 72_P67A-K31A* (KK2681), *hfq_1-72_V68A-K31A* (KK2687), *hfq_1-72_S69A-K31A* (KK2688), *hfq_1-72_H70A-K31A* (KK2682), *hfq_1-72_H71A-K31A* (KK2683) and *hfq_1-72_S72A-K31A* (KK2684)]. The experimental conditions were as for Figure 2B. Data are the average of three assays, two from one culture and one from an independent culture, and error bars represent the standard deviation of the mean. **(E)** β-galactosidase activity measured in isogenic strains carrying the P_BAD_-*chiP-lacZ* fusion, with R66A combined with various Hfq face mutants. Strains used: Δ*hfq* (DJS2689), WT (DJS2690), *hfqR66A* (KK2725), *hfqR16A* (DJS2693), *hfqR66A-R16A* (KK2726), *hfqK31A* (DJS2695), *hfqR66A-K31A* (KK2727), *hfq*_1-65_ *K31A* (KK04), *hfq_1-72_K31A* (KK2658) and *hfq_1-72_R66A-K31A* (KK2680). Strains were grown and assayed for β-galactosidase activity as in Figure 2B. Data are the average of three assays, two from one culture and one from an independent culture, and error bars represent the standard deviation of the mean. **(F)**. β-galactosidase activity measured in P_BAD_-*chiP-lacZ* strains with the Hfq acidic tip of CTD mutated to basic residues (Santiago-Frangos et al., 2017) or the linker region of the CTD deleted in combination with face mutants [Δ*hfq* (DJS2689), WT (DJS2690), *hfq_1-65_* (KK01), *hfq_Δlink_* (KK2558), *hfq_tipmut_* (KK2570), *hfq-R16A* (DJS2693), *hfq_1-65_R16A* (KK02), *hfq_Δlink_R16A* (KK2654), *hfq _tipmut_R16A* (KK2651), *hfqK31A* (DJS2695) and *hfq_1-65_K31A* (KK04), *hfq_Δlink_K31A* (KK2655) and *hfq _tipmut_K31A* (KK2652)] using the experimental conditions as in Figure 2B. Data are the average of three assays, two from one culture and one from an independent culture, and error bars represent the standard deviation of the mean.

The truncations of the Hfq CTD (Hfq_1-92_, Hfq_1-82_, Hfq_1-72_) were qualitatively compared to Hfq_1-65_ for regulation of *chiP-lacZ* on a lactose MacConkey plate (Supplementary Figure S5). WT colonies were white (full repression) while Δ*hfq* colonies were red, indicative of a lack of repression by ChiX. Hfq_1-65_ colonies had a pale red phenotype, consistent with modest derepression (Supplementary Figure S5B, compare to quantitation in Figure 2B). However, the colonies for each of the intermediate Hfq truncations were white, suggesting that only the region between residues 65 and 72 was necessary for full ChiX repression of *chiP-lacZ* by this assay. We note that this region was identified as involved in core packing, and is relatively conserved, compared to the rest of the CTD (41).

Given the apparent full function for Hfq_1-72_ in this assay, the Hfq_1-72_ truncation was further compared to Hfq_1-65_, alone or in combination with mutations on the RNA binding faces. Effective repression of *chiP-lacZ* was observed not only for Hfq_1-72_, but for Hfq_1-72_K31A, a very different result from the loss of repression in Hfq_1-65_K31A (Figure 3B). Therefore, Hfq_1-72_ also is fully functional by this more stringent assay. Combinations of Hfq_1-72_ with binding face mutants were also assayed for the sRNA-independent *mutS-lacZ* reporter (Figure 3C). As for *chiP*, Hfq_1-72_ and Hfq_1-72_K31A were both fully functional. Therefore, for the distal site defects observed in Hfq_1-65_ in combination with K31A, the region between amino acids 65-72 is critical, but the rest of the Hfq CTD, including the tip and linker regions, is not.

To identify the critical residue(s) in the region between residues 65 and 72 of the CTD, individual alanine substitutions were created between amino acids 66 and 72 in the context of the Hfq_1-72_K31A protein, and were tested for *chiP-lacZ* activity (Figure 3D). Mutation of R66, the most conserved amino acid in this region of Hfq, to alanine had the strongest effect, leading, in combination with K31A, to loss of repression of *chiP-lacZ*, although not quite as much as for Hfq_1-65_K31A. Other alanine mutations in this region combined with K31A gave repression similar to K31A itself, suggesting that they do not contribute significantly to function, at least individually, for this assay (Figure 3D). Based on these results, the synergism between the Hfq CTD and the distal site, reflected in the loss of regulation of two reporters, the *chiP-lacZ* reporter and the *mutS-lacZ* reporter, is likely due to sequences just beyond amino acid 65, particularly R66.

We also assayed another set of constructs with R66A in full-length Hfq (rather than the Hfq_1-72_ used in Figure 3D), in otherwise WT Hfq or combined with the R16A (rim) or K31A (distal) face mutants. Disruption of R66A by itself (Figure 3E) did not lead to a significant defect in *chiP*-*lacZ* regulation by ChiX, but in combination with K31A had a phenotype similar to if not quite as strong as that of Hfq_1-65_K31A, fully supporting the idea that R66 is the primary CTD residue that impacts distal site function.

Given that a truncation of Hfq to 72 amino acids was fully functional for ChiX regulation of *chiP*, we would not expect to see a defect in regulation in cells carrying mutation of the Hfq acidic tip or the linker region, just N-terminal of the tip. As expected, the tip RNKN mutant (called here Hfq_tipmut_) was fully functional for ChiX repression of *chiP*-*lacZ* (compare to Hfq_1-65_ and WT), and retained function in combination with K31A (Figure 3F). The Δ*link* mutation also was functional in combination with K31A. Thus, for this assay, in agreement with the result with truncations, the linker and tip do not appear to play significant roles.

### Additive effect of CTD deletion and rim mutations for Class I sRNA-dependent regulation

For Class I sRNAs, the sRNA binds to the proximal and rim sites of Hfq, and targets bind to the distal face. We next examined the effect of the Hfq_1-65_ mutation and combinations of Hfq mutants on repression of a *sodB-lacZ* fusion by the Class I sRNA RyhB. This assay was carried out either in cells deleted for the Fur repressor (Figure 4A) or in cells in which RyhB was overexpressed from a plasmid (Figure 4B). In the absence of Fur, RyhB is expressed at high levels from the chromosomal copy of the gene; however, this level of RyhB is less than that achieved with induction of a multicopy plasmid. This is most obvious in comparing the effects of the *hfqQ8A* or *hfq_1-65_Q8A* mutations to the deletion of *hfq*. HfqQ8A significantly disrupts binding of all sRNAs to the proximal face, leading to instability of the sRNAs (8) (38). In the Δ*fur* strain, this results in full loss of regulation; in the pBR-RyhB strain, HfqQ8A reduces but does not eliminate regulation, presumably because higher RyhB expression can partially overcome the instability of the sRNA. We note that in this case, Hfq_1-65_ exacerbates the defect in the Q8A mutant (Figure 4B), though the Hfq_1-65_ mutation, on its own, somewhat reduced the effectiveness of RyhB repression.

**Figure 4.**
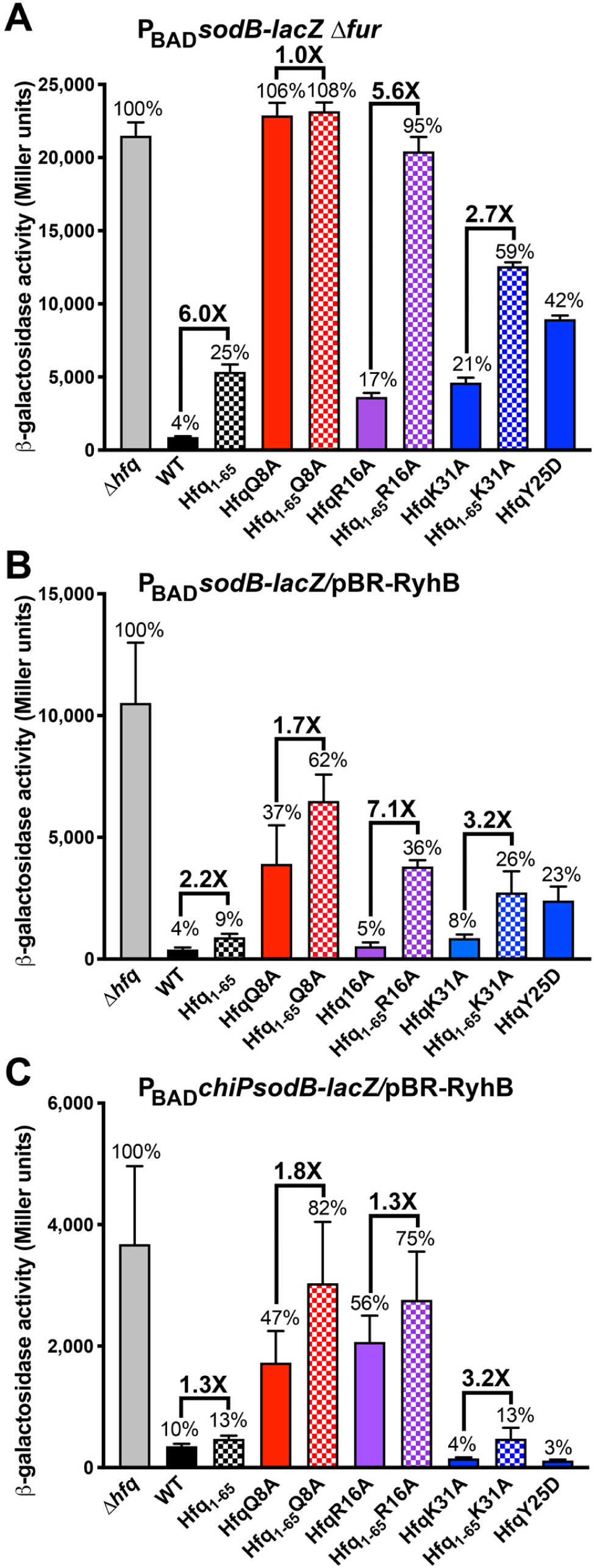
Effect of the Hfq CTD in regulation by Class I sRNA RyhB. (**A**) β-galactosidase activity measured in isogenic derivatives all carrying P_BAD_*-sodB-lacZ Δfur::zeo ryhB*^+^; RyhB is expressed at a high level from the chromosome because the Fur repressor is absent. *hfq* mutants were at the native locus [Δ*hfq* (KK2706), WT (KK2693), *hfq_1-65_* (KK2694), *hfqQ8A* (KK2695), *hfq_1-65_Q8A* (KK2717r*), *hfqR16A* (KK2696), *hfq_1-65_R16A* (KK2707), *hfqY25D* (KK2719); *hfqK31A* (KK2701), *hfq_1-65_K31A* (KK2702) and *hfqY25D* (KK2719)]. Cells were grown in flasks in LB with .002% arabinose to OD_600_ ∼ 1.0. (**B**) These strains carry the same fusion as in A, but are *fur*^+^; RyhB is expressed from the *lac* promoter on a pBR-RyhB plasmid. Strains were grown in LB medium with 100 μg/ml ampicillin, 10 μM IPTG, and 0.002% arabinose at 37°C to OD_600_ ∼ 1.0 and assayed for β-galactosidase activity in isogenic derivatives of DJS2676 carrying the P_BAD_*-sodB-lacZ* fusion [Δ*hfq* (DJS2682. WT (DJS683), *hfq_1-65_* (KK2611), *hfqQ8A* (DJS2684), *hfq_1-65_Q8A* (KK2613r*), *hfq16A* (DJS2686), *hfq_1-65_R16A* (KK2615); *hfqK31A* (DJS2688), *hfq_1-65_K31A* (KK2619) and *hfqY25D* (DJS2687)], all carrying a plasmid expressing RyhB (pBR-RyhB). Data are from triplicate independent cultures. (**C**) β-galactosidase activity measured in a reporter in which the UA-rich Hfq binding region of the *chiP* mRNA is fused to the RyhB base pairing region of *sodB* (8), grown and assayed as for Figure 4B. Strains are all *hfq* mutant derivatives of P_BAD_*-chiP-sodB-lacZ* (DJS2985), [Δ*hfq* (DJS3007), WT ( DJS3009), *hfq_1-65_* (KK2622), *hfqQ8A* (KK2623), *hfq_1-65_Q8A* (KK2624r*), *hfqR16A* (DJS3010), *hfq_1-65_R16A* (KK2626), *hfqK31A* (KK2629), *hfq_1-65_K31A* (KK2630) and *hfqY25D* (DJS3011)] carrying plasmid pBR-RyhB. All assays were done with three biological replicates, and error bars represent the standard deviation. Note that all strains used in Figures 4A and 5B-D were assayed together, and thus the same values appear for the same strains in the different panels.

Consistent with previous results, RyhB repression of the *sodB-lacZ* fusion was only modestly disrupted by mutation of R16 at the rim or K31A at the distal site; Y25D at the distal site was somewhat more defective (38) (Figures 4A and 4B). However, unlike the results for

ChiX and *mutS* discussed above, the combination of K31A and Hfq_1-65_, while more defective than either K31A or Hfq_1-65_, was not fully defective (Figure 4A and 4B). *sodB* and other Class I targets bind to the distal face of Hfq; apparently this binding, while somewhat sensitive to loss of the CTD in a weak distal site mutant is not nearly as sensitive to the combination of K31A and the CTD deletion as seen for repression of *chiP* and *mutS* (Figure 2).

Consistent with a role for the rim in pairing and RyhB stability, the combination of rim mutant R16A with Hfq_1-65_ was significantly more defective than either one alone (Figure 4A). Unlike the effect of ChiX overexpression for *chiP-lacZ* regulation (Figure S4) however, the Hfq_1-65_R16A defect was still significant when RyhB was further overexpressed (Figure 4B).

In previous work, we created a chimeric reporter for RyhB, *chiP-sodB-lacZ*, in which the predicted rim Hfq binding site of *chiP* replaced the distal Hfq binding site of *sodB* in an mRNA containing the pairing region of *sodB*. The resulting fusion is regulated by RyhB, and, unlike the original *sodB* target, this chimeric fusion is defective for RyhB repression in rim mutants and not defective in distal site mutants (8). Because regulation of this fusion should be independent of distal site defects, we expected the fusion would be less sensitive to the distal site defects in the Hfq_1-65_ truncation. Consistent with previous results (8), mutations in the distal site (Y25D and K31A) allowed very effective repression of the chimeric reporter (better than that seen for the WT construct), possibly because RyhB, unable to pair with its usual targets, is stabilized and available for pairing with this target (Figure 4C). Hfq_1-65_ modestly perturbed the tight repression seen in the K31A mutant (Figure 4C). The repression of *chiP*-*sodB* by the Hfq_1-65_K31A double mutant protein confirms that this Hfq derivative must be functional; total loss of Hfq (Δ*hfq*) abrogates regulation. For this fusion, however, rim mutations were defective in the context of either full length Hfq or Hfq_1-65_ (Figure 4C).

### Hfq CTD requirements for RyhB differ from those for ChiX

To further define the CTD requirements for RyhB-dependent repression of *sodB-lacZ*, we examined repression in several of the same *hfq* mutants tested for ChiX repression of *chiP*-*lacZ* (Figure 5A). Unlike ChiX repression, where Hfq_1-72_ had little or no effect, Hfq_1-72_ reduced repression by RyhB 3.3-fold relative to WT, and had a similar 2.7-fold defect relative to R16A (Figure 5B). However, Hfq_1-72_K31A was only slightly less effective than K31A alone (1.1 fold, Figure 5B). Therefore, for RyhB repression of *sodB*, Hfq_1-72_ behaves similarly to full-length Hfq for distal site function (binding *sodB* mRNA in this case), but is defective for rim function.

**Figure 5.**
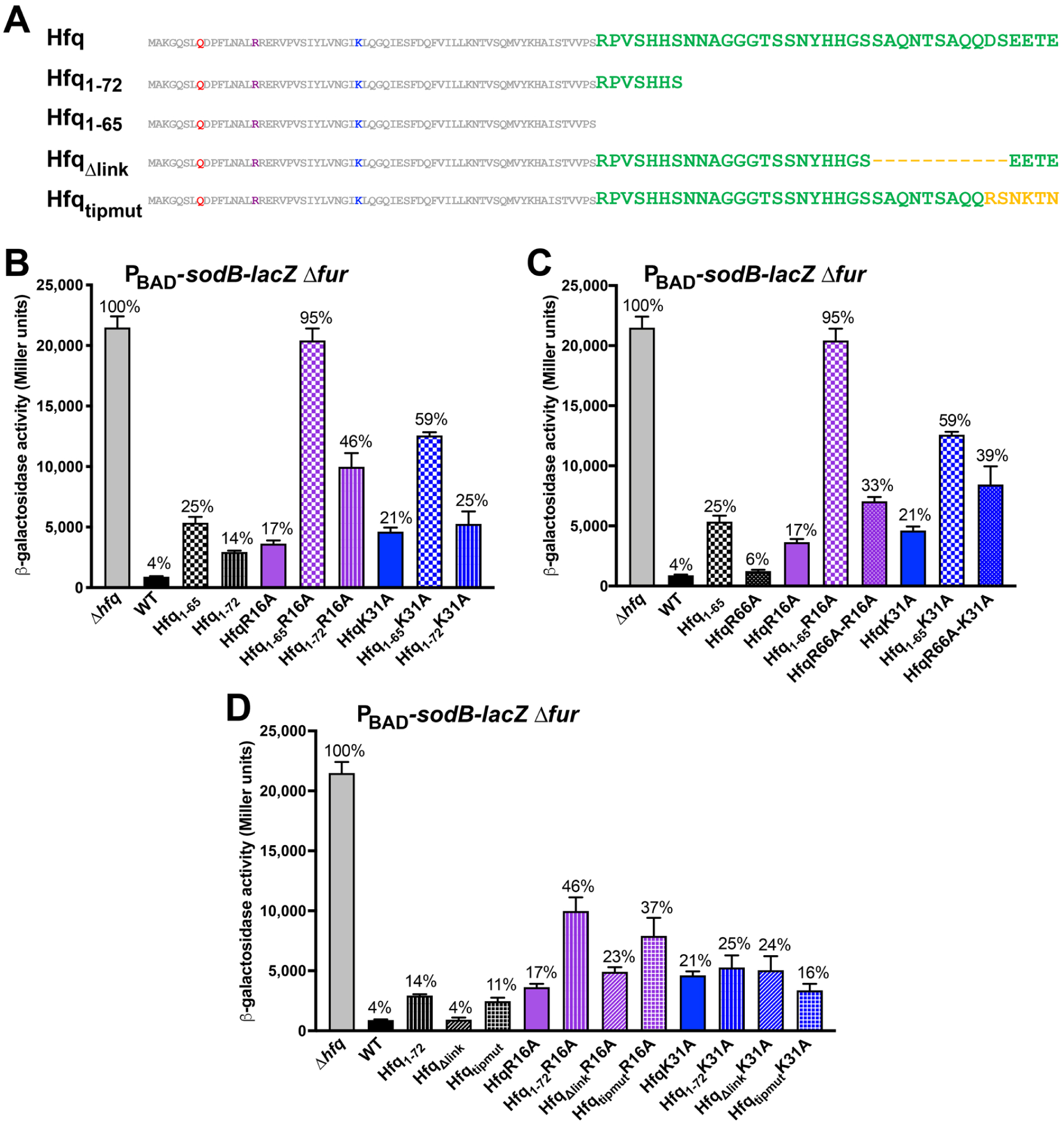
Dissecting the Hfq CTD for regulation by a Class I RNA. (**A**) Schematic of Hfq CTD truncations and mutations. (**B**) β-galactosidase activity measured in isogenic strains of P_BAD_*- sodB-lacZ ΔfurA::zeo* expressing Hfq mutants [*Δhfq* (KK2706), WT (KK2693), *hfq_1-65_* (KK2694), *hfq_1-72_* (KK2718), *hfqR16A* (KK2696), *hfq_1-65_R16A* (KK2707), *hfq_1-72_R16A* (KK2704), *hfqK31A* (KK2701), *hfq_1-65_K31A* (KK2702) and *hfq_1-72_K31A* (KK2705)], which were grown as for Figure 4A. (**C**) β-galactosidase activity measured in isogenic strains of P_BAD_*-sodB-lacZ ΔfurA::zeo* expressing Hfq mutants [*Δhfq* (KK2706), WT (KK2693), *hfq_1-65_* (KK2694), *hfq*R66A (KK2731), *hfqR16A* (KK2696), *hfq_1-65_R16A* (KK2707), *hfq*R66AR16A (KK2732r*), *hfqK31A* (KK2701), *hfq_1-65_K31A* (KK2702) and *hfq*K31AR66A (KK2733r*)]. (**D**) β-galactosidase activity measured in isogenic strains of P_BAD_*-sodB-lacZ ΔfurA::zeo* expressing Hfq mutants [*Δhfq* (KK2706), WT (KK2693), *hfq_1-72_* (KK2718), *hfq_Δlink_* (KK2699), *hfq_tipmut_* (KK2700), *hfqR16A* (KK2696), *hfq_1-72_R16A* (KK2704), *hfq_Δlink_R16A* (KK2697), *hfq_tipmut_R16A* (KK2698), *hfqK31A* (KK2701), *hfq_1-72_K31A* (KK2705), *hfq_Δlink_K31A* (KK2715), and *hfq_tipmut_K31A* (KK2714)], which were grown in LB with 0.002% arabinose. The experimental conditions were as for Figure 4A. Note that experiments shown in Figures 4A and 5 were assayed together, and the same values are thus used for various samples found in multiple panels. All assays were done with three biological replicates, and error bars represent the standard deviation.

For ChiX regulation of *chiP-lacZ*, R66 was a critical residue, leading to a significant defect in combination with K31A (Figure 3E). For RyhB-mediated repression of *sodB-lacZ*, R66A had a modest effect by itself (1.4-fold), and a slightly more significant defect in combination with R16A (1.9-fold) or K31A (1.8-fold) (Figure 5C).

Given the requirement for sequences beyond amino acid 72 in the Hfq CTD for RyhB regulation of *sodB* (Figure 5B), we compared the effect of mutations in the acidic tip and linker to the 1-72 truncation (Figure 5D). Unlike the results with ChiX repression of *chiP* (Figure 3F), RyhB repression of *sodB* was sensitive to mutations in the tip, both in an otherwise WT strain (2.7-fold) or in the context of the R16A rim mutant (2.2-fold), but not in the K31A derivative (0.7-fold) (Figure 5D). The Hfq_Δlink_ mutant was functional on its own and in combination with R16A or K31A (1.0-fold, 1.4-fold, and 1.1-fold, respectively). These results support the idea that the tip is needed for full rim function of Hfq, while mutation of R66 impacts both rim and distal face function.

### Class II sRNAs show varied accumulation in CTD mutant backgrounds

Given that defects in regulation sometimes correlate with differences in sRNA levels, we next selected a set of Class II and Class I sRNAs and examined their accumulation in the various Hfq mutant backgrounds compared to the WT and Δ*hfq* strains.

We had previously found that Class II sRNAs ChiX and MgrR were more unstable in the Hfq_1-65_ strain and that all of the measured Class II sRNAs accumulated to lower levels in Hfq_1-65_, presumably due to the lower stability (18). Here we confirmed the lower accumulation of Class II sRNAs ChiX, MgrR and CyaR in Hfq_1-65_ (Figure 6A). This effect of Hfq_1-65_ on Class II sRNA levels was also observed in combination with both the R16A mutation and the K31A mutation. The low levels in Hfq_1-65_K31A are consistent with the lack of distal face function observed above (Figure 3B, C). We note that mutation of R66A, while partially limiting ChiX activity (Figure 3D, 3E), did not reduce levels of the Class II sRNAs significantly (Figure 6A, lane 13 compared to 12, and lane 19 compared to 18)

**Figure 6.**
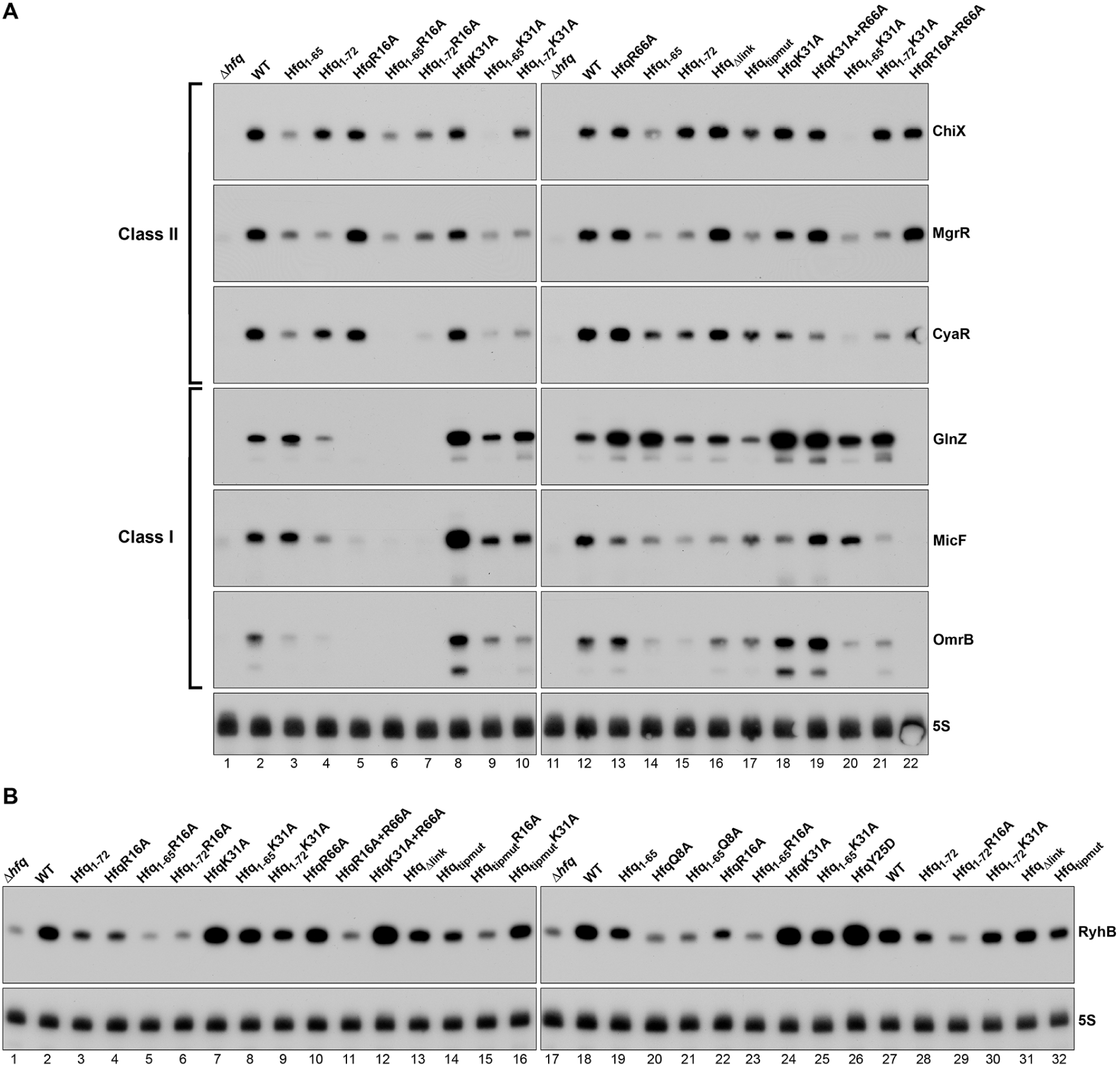
Effect of single and double *hfq* mutants on sRNA levels. (**A**) ChiZ, MgrR, CyaR, GlnZ, MicF and OmrB sRNAs levels in strains containing the *chiP-lacZ* fusion and grown in LB as in Figure 3 to OD_600_ ∼ 1.0. Strains used are those from Figures 2 and 3: [*Δhfq* (DJS2689; lane 1, 11), *hfq*^+^ (DJS2690; lanes 2, 12), *hfq_1-65_* (KK01; lanes 3, 14), *hfq_1-72_* (KK2438, lanes 4, 15), *hfq*R16A (DJS2693, lane 5), *hfq_1-65_R16A* (KK02, lane 6), *hfq_1-72_R16A* (KK2657, lane 7), *hfqK31A* (DJS2695, lanes 8, 18), *hfq_1-65_K31A* (KK04, lanes 9, 20), *hfq_1-72_K31A* (KK2658, lanes 10, 21), *hfq*R66A (KK2725, lane 13), *hfq_Δlink_* (KK2558, lane 16), *hfq_tipmut_* (KK2570, lane 17), *hfqK31AR66A* (KK2680, lane 19), *hfqR16AR66A* (KK2726, lane 22).] (**B**) RyhB levels in Δ*fur sodB-lacZ* fusion derivatives. Strains and growth conditions were as in Figure 4A. Strains used are the same or the original versions of remade strains (Table S1), as in Figures 4A and 5: [*Δhfq* (KK2706, lanes 1, 17), *hfq*^+^ (KK2693, lanes 2, 18), *hfq_1-72_* (KK2718, lanes 3, 28), *hfq*R16A (KK2696, lanes 4, 22), *hfq_1-65_R16A* (KK2707, lanes 5, 23), *hfq_1-72_R16A* (KK2704, lanes 6, 29), *hfqK31A* (KK2701, lanes 7, 24), *hfq_1-65_K31A* (KK2702, lanes 8, 25), *hfq_1-72_K31A*, (KK2705, lanes 9, 30), *hfqR66A* (KK2731, lane 10), *hfqR16AR66A* (KK2732, lane 11), *hfqK31AR66A* (KK2733, lane 12), *hfq_Δlink_* (KK2699, lanes 13, 31), *hfq_tipmut_* (KK2700, lanes 14, 32), *hfq_tipmut_R16A* (KK2698, lane 15), *hfq_tipmut_K31A* (KK2714, lane 16), *hfq_1-65_* (KK2694, lane 19), *hfqQ8A* (KK2695, lane 20), *hfq_1-65_Q8A* (KK2717, lane 21), *hfqY25D* (KK2719, lane 26)]. For both A and B, total RNA was subject to northern analysis using ^32^P-labeled oligonucleotide probes complementary to the indicated sRNAs and 5S (RNAs were probed sequentially on the same membrane).

Interestingly, however, accumulation of the Class II sRNAs varied in other C-terminal mutants, with each Class II sRNA having a somewhat distinct pattern. For ChiX, in which the Hfq_1-72_ derivative did not affect regulation (Figure 3B), this truncation had very modest effects on total levels, either in an otherwise wild-type Hfq or in the K31A mutant (Figure 6A, lanes 4 and 10 compared to lane 2 and 8, respectively). It did reduce levels of ChiX somewhat in combination with R16A (lane 7). Deletion of the Hfq tip (Hfq_tipmut_) had a modest effect on ChiX levels (lane 17) and the deletion of the link (Hfq_Δlink_) had no effect (lane 16), consistent with only a minor effect of these alleles on regulation by ChiX. In contrast, the levels of MgrR were similarly low for Hfq_1-65_, Hfq_1-72_ and Hfq_tipmut_ in otherwise WT Hfq (Figure 6A, lanes 3, 4, 17). These effects were not significantly exacerbated by R16A or K31A (Figure 6A, lanes 6, 7, 9, 10). CyaR levels were only slightly lower in the Hfq_1-72_ and Hfq_tipmut_ backgrounds (Figure 6A, lanes 4, 17) but, unlike what was observed for MgrR, the combination of Hfq_1-72_ with either R16A or K31A led to even lower levels (Figure 6A, lanes 6, 7, 9, 10).

### Extent of Class I sRNA accumulation in CTD mutant backgrounds is sRNA specific

For Class I sRNAs, we chose to examine the levels of three sRNAs showing different effects of Hfq_1-65_ relative to WT in the RNA-Seq experiments: GlnZ, the only sRNA showing increased levels, MicF, showing modestly decreased levels, and OmrB, with significantly decreased levels and previously shown to have reduced intrinsic stability in Hfq_1-65_ (18) (Figure 1, Table S4, Supplementary Figure S2B). These sRNAs were probed on the same membrane used for ChiX, MgrR and CyaR (Figure 6A). While GlnZ levels were elevated in both the Hfq_1-65_ (compare lane 3 to 2 and lane 14 to 12) and HfqR66A (compare lane 13 to 12) backgrounds, the effects of these mutations on MicF and OmrB levels were mixed. Hfq_1-65_ was associated with reduced OmrB while there was less of an effect on MicF levels (compare lanes 3 to 2 and lanes 14 to 12), and HfqR66A was associated with reduced MicF levels with less of an effect on OmrB levels (compare lane 13 to 12). In contrast, the levels of all three sRNAs generally were low in Hfq_1-72_, Hfq_Δlink_ and Hfq_tipmut_, and almost undetectable in all HfqR16A mutants (compare lanes 4-7 to lane 2 and lane 15-17 and 22 to lane 12). As expected for Class I sRNAs (8), the levels of GlnZ, MicF (in one sample), and OmrB were higher in the HfqK31A background compared to WT (compare lanes 8 to 2 and 18 to 12). This accumulation was decreased for HfqK31A in combination with Hfq_1-65_ and Hfq_1-72_, but not for the HfqK31A+R66A combination.

We additionally examined RyhB in the Δ*fur* strains used to measure RyhB-dependent repression of *sodB-lacZ* fusions (Figure 6B); these strains have elevated levels of RyhB due to the absence of the Fur repressor. RyhB levels were lower but still detectable in the absence of Hfq (lanes 1 and 17 compared to lanes 2 and 18). Compared to the WT strain, the Hfq_1-65_ and Hfq_1-72_ derivatives had somewhat lower RyhB levels (lane 3 compared to 2, lane 19 compared to 18). Hfq binding, particularly on the critical proximal face, is required for stabilization of Class I sRNAs. Consistent with this, RyhB levels in strains expressing HfqQ8A were as low as in the absence of Hfq (compare lanes 20, 21 to lane 17), and were reduced in HfqR16A (compare lane 4 to lane 2 and lane 22 to lane 18). In the R16A background, the Hfq_1-65_, Hfq_1-72_ and Hfq_tipmut_ mutations all further lowered RyhB levels (lanes 5, 6, 15 compared to lane 4). These reductions in RyhB levels correlate well with reductions in repression (Figures 4A, 5B, 5D) and are consistent with our hypothesis that the CTD region beyond amino acid 72, and in particular the acidic tip, is required for Hfq rim binding to RyhB and intrinsic RyhB stability. Similar to GlnZ and OmrB, RyhB levels were higher in HfqK31A as well as in the HfqY25D background compared to WT cells (compare lane 7 to lane 2, compare lanes 24 and 26 to lane 18 and 27). In both the WT and K31A backgrounds, the R66A mutation had no effect on either function (Figure 5C) or RyhB levels (compare lane 10 to lane 2, lane 12 to lane 7).

Overall, the different effects of Hfq_1-65_ and Hfq_1-72_ on the levels of Class I sRNAs are again consistent with multiple roles of the CTD. Disrupting or deleting R66A has a profound effect on GlnZ, increasing its levels, similar to a distal face defect, further supporting the importance of R66 for the distal face. For other sRNAs such as RyhB, only the region beyond amino acid 72 impacts stability. These differences in the effects of the mutations on GlnZ, MicF, OmrB and RyhB levels also reinforce the idea that each individual sRNA has somewhat different interactions with Hfq.

## DISCUSSION

The role of the *E. coli* Hfq CTD has been puzzling. The conserved core of the protein, amino acids 1-65, is present in most structures and includes three RNA binding faces, as shown in Figure 2A. Amino acids 66-102 are less conserved and have not been visible in most structures, suggesting they may be disordered. *In vitro* studies have implicated the acidic tip of *E. coli* Hfq in decreasing non-specific RNA binding and promoting the release of dsRNA after pairing, likely via interactions with the basic patches at the rim of Hfq (17). However, different groups have reported rather varied conclusions in terms of the need for the CTD for regulation by sRNAs *in vivo* (summarized in Table S6, and discussed further below). Here, we used chromosomally-encoded Hfq variants and a range of quantitative and qualitative assays to further explore the role of the CTD *in vivo*. Our results point to at least two distinct roles for the region of Hfq beyond amino acid 65. In both cases, the *in vivo* consequence of disrupting these roles are modest unless one of the RNA binding faces of the conserved core is also disrupted by mutation. These results suggest that the Hfq CTD acts in a redundant manner *in vivo*, reinforcing and optimizing the roles of the Hfq RNA binding faces in RNA trafficking and regulation. This redundancy as well as the bipartite roles of the CTD provide some explanation for the range of previous conclusions about whether the CTD is in fact critical for regulation.

### Role for arginine 66 in Hfq distal face function

The first role of the CTD is defined by the difference in behavior of a truncation to residue 65 compared to truncation to residue 72. Hfq_1-65_, while only modestly defective in our assays, had a significant effect when combined with mutations in the rim or distal binding faces of Hfq (Figure 2B, C, Figure 4A). Given that Hfq_1-72_ was not as defective for assays dependent on the Hfq distal face and was not synergistic with the K31A mutation for these assays (Figure 3B, C), the critical region for this distal site activity must be between amino acids 65 and 72. Our analysis identified R66, the best-conserved amino acid in this region, as the most critical for this distal face activity (Figure 3D, E). R66A, with the rest of the CTD intact, partially mimicked the loss of the full CTD for regulatory activity when the weak distal face mutation K31A was also present (Figure 3E and 5C).

The R66A single mutant also had increased levels of the Class I sRNA GlnZ, again mimicking the effect of deleting the full CTD (Figure 6). Because Class I sRNAs are usually displaced from Hfq after pairing, and displacement allows them to be rapidly degraded, less pairing increases stability and thus accumulation of these sRNAs (8). The GlnZ sRNA might be particularly sensitive to degradation with use or its targets may be particularly vulnerable to any distal site disruption, so that a disruption in GlnZ target binding in the Hfq_1-65_ or R66A mutants results in increased accumulation.

Perhaps the region just beyond amino acid 65, and in particular the well-conserved amino acid R66, should be considered a transition between the conserved core of Hfq and the disordered extended CTD, with a role in the full activity of the distal face of Hfq. This region of the CTD is less disordered and packs against the Hfq core in some structures (13,17,41,42). We note, however, that R66A did not compromise the stability of the Class II sRNAs, suggesting that the more extended region around R66 up to amino acid 72 contributes to Class II binding to the distal site, even though the mutation in R66 alone is sufficient to compromise activity.

### Role for the CTD tip in Hfq rim function

A second and distinct role of the CTD is played by the region beyond amino acid 72, particularly the tip of the Hfq CTD. This part of the CTD is disordered (41). Recent *in vitro* studies on the acidic tip suggested that this tip is necessary for the properties attributed to the CTD (17). In that study, *de novo* Rosetta modeling and competitive binding experiments showed that the acidic tip of CTD transiently binds to the basic arginine patch on the rim of the Hfq core. The data suggest a model in which the acidic tip mimics a nucleic acid, reducing nonspecific RNA binding and supporting release of dsRNA, the expected product of pairing of sRNA and mRNA.

Hfq derivatives Hfq_1-72_, Hfq_Δlink_ and Hfq_tipmut_, alone and in combination with the rim mutants, support a role for the CTD beyond amino acid 72 in mediating events at the rim of Hfq, consistent with the *in vitro* data. Hfq_1-72_R16A and Hfq_tipmut_R16A were significantly more defective for RyhB regulation of *sodB-lacZ* than either mutation alone (Figure 5B, D). These results suggest that, unlike the role of R66 and the region between amino acids 65 and 72, the extended CTD (particularly the tip) affect Hfq functions associated with the rim of Hfq. We cannot distinguish between the tip region affecting Class I sRNA intrinsic stability or other functions such as pairing and release; the CTD may collaborate with the rim to affect all of these.

We note that the Hfq_1-72_ mutant, while not affecting distal-site regulation by ChiX (Figure 3B), does lead to lower levels of some Class II sRNAs (Figure 6). While we do not have a full explanation for this, the change in Class II sRNAs levels may reflect either decreased binding of Class II targets to the rim or more effective pairing; each may promote faster displacement from Hfq and degradation after pairing. Thus, loss of the CTD tip, which could increase the binding of some Class II targets to the rim, may allow more rapid turnover without loss of regulation.

### Integrating current observations with previous findings

Based on our findings, we revisited the published *in vivo* studies on the Hfq CTD in *E. coli* to see if our analysis can help in interpreting the apparent contradictions. *In vitro* studies do support the idea of two functionally distinct regions of the *E. coli* Hfq CTD, with amino acids 66-70 at the very beginning of the Hfq CTD more ordered and interacting with the distal face of Hfq (41, 42) and residues beyond amino acid 70 disordered and showing evidence of interactions with the rim of Hfq (41). We note that the *in vivo* phenotypes are not as strong as those observed *in vitro*. A possible explanation for this discrepancy is that the kinetic parameters that drive the *in vitro* observations may be less important *in vivo* where there is a large pool of different RNAs that compete for binding to the various Hfq surfaces.

In Supplementary Table S6, we summarize the *in vivo* studies of the *E. coli* Hfq CTD, the assays used to evaluate Hfq function, and how the results compare to what we show here. We present these in chronological order to reflect the increasing knowledge of the cellular roles of Hfq, resulting in changes in what was assayed. In all but the very first previous studies, the *hfq* mutant alleles were expressed from plasmids, and in many, Hfq was overexpressed, further complicating comparisons to the current work. Double mutant derivatives such as those used here were not examined in any of the previous studies.

Overall, given the modest effects we saw on downstream mRNA levels in our RNA-Seq experiments, it is not surprising that many studies found no critical requirement for the Hfq CTD for regulation (15, 40) (Sonnletiner et al, Olsen et al, Table S6). However, two studies had significant inconsistencies with our results (16, 41), reporting that Hfq_1-65_ failed to support many Hfq-dependent effects, including expression of RpoS and RyhB regulation of *sodB-lacZ* (Vecerek et al, Beich-Fransen et al, Table S6). In both of these studies, Hfq was overexpressed from the same plasmid without a description of the Hfq levels. It is possible that Hfq_1-65_ could aggregate or otherwise become inactive under these overproduction conditions. For other studies where Hfq-mediated regulation showed some dependence on the near-core region of the Hfq CTD, we suspect the assays were particularly dependent on strong or longer-term binding of mRNAs to the distal face of Hfq. These tests include autoregulation of Hfq (43) and positive regulation of GlmS translation (44) (Caillet et al and Salim et al, Table S6), and are consistent with the effects we observed for Hfq repression of *mutS* (Figure 2C).

In other studies, levels of sRNAs were examined, and modest differences of the sort we detected were observed. For instance, Olsen et al. found lower levels of ChiX in strains expressing Hfq_1-65_ or Hfq_1-66_, but not in those expressing Hfq_1-72_; the decrease in ChiX was not sufficient to lead to any significant defect in regulation in functional assays (40). One clear conclusion from our current and past experiments is that a decrease in sRNA levels, while diagnostic of some aspects of Hfq function, is not always correlated with changes in regulatory activity (8). For instance, Class I sRNAs are more stable and thus accumulate to higher levels in distal site mutants that slow or block regulation (8); we see this in the present study as well, with higher Class I sRNAs in the HfqK31A distal face mutant although regulation is defective (Figures 4, 6). In other cases, we do not fully understand the basis for the effects on sRNA levels. For instance, while a strain carrying HfqK31AR66A was almost as defective for *chiP* regulation as one expressing Hfq_1-65_K31A, the ChiX levels are very low for Hfq_1-65_K31A and significantly higher for HfqK31AR66A (Figures 3E and 6A), Thus sRNA levels on their own cannot be used to predict or infer function.

While we conclude here that removing the Hfq CTD, in otherwise WT Hfq, has only modest effects on regulation, it is still possible that there are more significant consequences for deleting the Hfq CTD under changing growth conditions or in the kinetics of the response as sRNAs are induced or start to disappear from the cell. We also have not examined the role of the Hfq CTD on other Hfq features such as subcellular localization, binding to DNA or binding to other proteins important for RNA metabolism (17,45,46). It will be interesting to determine how the differential effects of Hfq CTD mutations on various sRNAs reflect these other features as well as how the differential effects impact the competition among RNAs for Hfq binding.

In general, the differences in how Class I and Class II sRNAs use Hfq, coupled with the two roles for the CTD discovered in our work and variation in the effects of different mutations on individual sRNAs, provide an explanation for why different assays by different groups have come to inconsistent conclusions. As discussed above, the accumulation for each Class I and Class II is different, emphasizing the variations in sRNA function. For Class I sRNAs, loss of the full CTD disrupts target binding to the distal face, leading to stabilization of some sRNAs, while simultaneously destabilizing the sRNAs due to loss of rim contacts. The balance between these effects differs for each sRNA (Figure 6). For Class II sRNAs, the disruption of the distal face generally leads to less sRNA accumulation, which is not compensated by disruption of the rim, though again the effects of different mutant combinations on levels varied among the sRNAs. Understanding the basis of these differences will require further *in vivo* and *in vitro* studies.

Overall, our results suggest that the CTD supports the roles previously defined for the Hfq faces. The R66 residue contributes to distal face function, while the extended tip of the disordered CTD has its most critical functions in connection with the rim. It is worth pointing out that, because of these roles of the CTD tip, C-terminal tags on Hfq may lead to modest defects in regulation, similar to loss of the extended CTD, and may be particularly deleterious in combination with Hfq binding face mutants. While we did not observe strong phenotypes for Hfq_1-65_ under the growth in rich media examined here, the CTD may become important *in vivo* under conditions that require optimal availability and kinetics of Hfq function. Finally, the differential effects we observed for various sRNAs underscore how the interaction of each sRNA with Hfq is unique.

## DATA AVAILABILITY

The NGS data discussed in this publication have been deposited in NCBI’s Gene Expression Omnibus (Edgar *et al*., 2002) and are accessible through GEO Series accession number GSE139988 (https://www.ncbi.nlm.nih.gov/geo/query/acc.cgi?acc=GSE139988).

## SUPPLEMENTARY DATA

Supplementary Data are available at NAR online.

## ACKNOWLEDGEMENT

We thank the NICHD Molecular Genomics Core for library sequencing and Konrad Förstner for valuable discussions on using READemption. We thank Daniel Schu for constructing many of the strains used in this work and for his comments and advice during the work. This work utilized the computational resources of the NIH HPC Biowulf cluster and was supported by the NIH intramural research programs of NCI and NICHD. We thank Sarah Woodson and members of the Gottesman and Storz laboratories for comments on the manuscript. We thank Ricardo Francis for construction of some of the strains.

