## Supplemental Material for "Multiple *in vivo* roles for the C-terminal domain of the RNA chaperone Hfq"

**List of Supplemental Files**

**Supplemental Figures:**

**Figure S1.** Reproducibility of RNA-Seq data for full length Hfq and $Hfq_{1-65}$.

**Figure S2.** Volcano plot for co-IP of RNAs with FLAG-tagged Hfq from WT and $Hfq_{1-65}$ and genome browser for selected RNAs.

**Figure S3.** Levels of Hfq single and double mutants characterized in study.

**Figure S4.** Multicopy ChiX can suppress the regulatory defect for ChiX regulation of *chiP-lacZ* in a $Hfq_{1-65}$ K31A background.

**Figure S5.** Effect of Hfq CTD truncations on ChiX-based regulation of *chiP-lacZ*.

**Supplemental Tables:**

**Table S1.** Strains used in study.

**Table S2.** Oligonucleotides used in study.

**Table S3.** RNA-Seq: levels of all RNA species in total RNA and IP samples for WT and $Hfq_{1-65}$.

**Table S4.** RNA-Seq: total RNA results, two-fold change or better, $Hfq_{1-65}$/WT.

**Table S5.** RNA-Seq: Hfq immunoprecipitation, two-fold change or better, $Hfq_{1-65}$/WT.

**Table S6:** Summary of Literature on *E. coli* Hfq CTD *in vivo*.

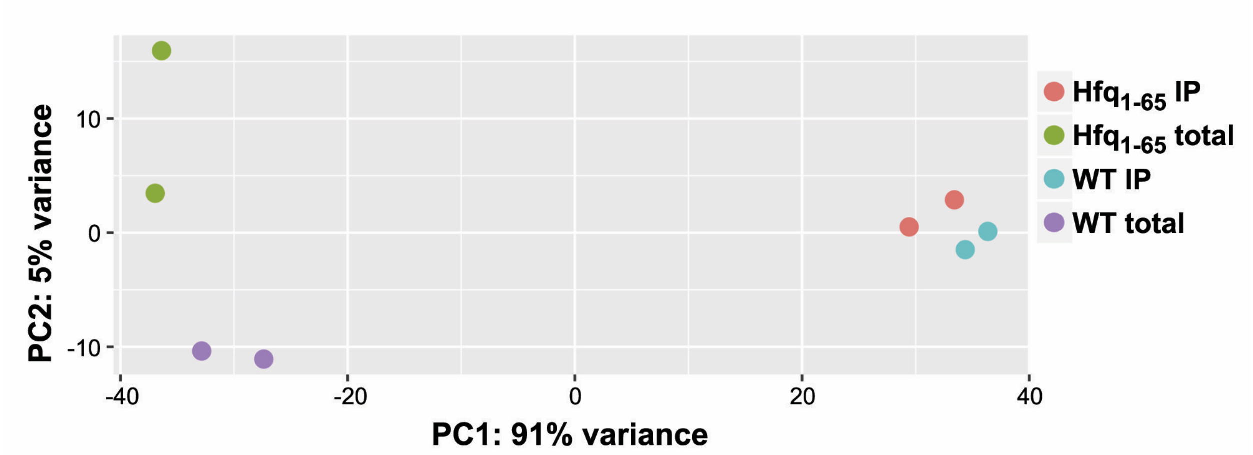


**Figure S1.** Reproducibility of RNA-Seq data for full length Hfq and $Hfq_{1\text{-}65}$. PCA plot for WT Hfq (KK2440) and $Hfq_{1\text{-}65}$ (KK2448) total and co-immunoprecipitation (co-IP) with Hfq RNA with biological replicates clustering together.

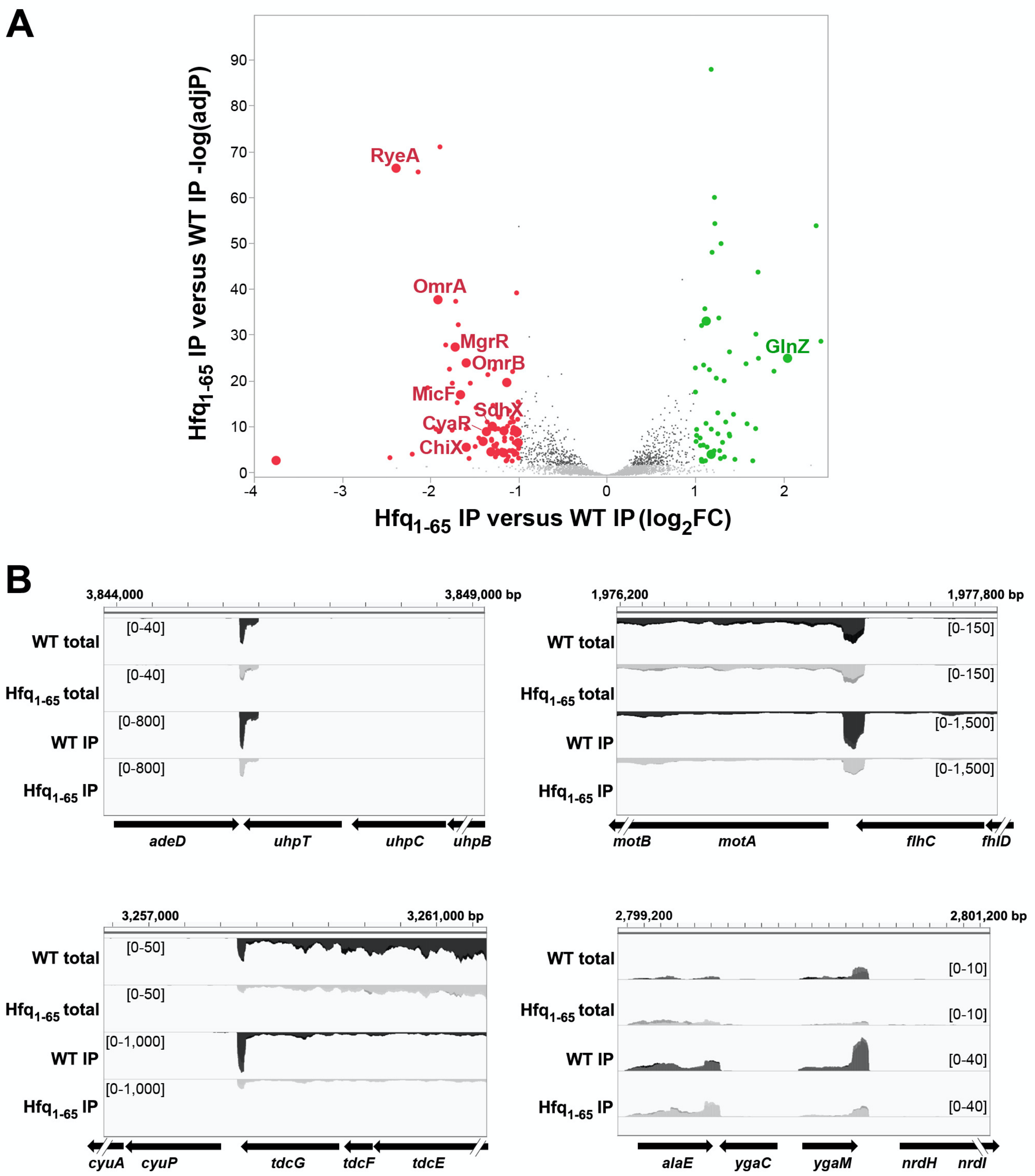


**Figure S2.** Volcano plot for co-IP of RNAs with anti-Hfq from WT and Hfq$_{1-65}$ and genome browser for selected RNAs. (**A**) Volcano plot comparing co-IP RNA from Hfq$_{1-65}$ (KK2448) to Hfq WT(KK2440). All genes with <2X change are shown in black or grey (signals too low to be significant). The red and green dots represent RNAs down and up regulated more than two-fold, respectively. A subset of sRNAs are labeled. (**B**) Genome browser images of data from total and Hfq co-IP RNA-Seq, showing patterns for two recently identified new sRNAs (UhpU, at the 3' UTR of *uhpT*, and MotR, at the 5' UTR of *motA* (1); unpublished, Storz lab), as well as two regions with patterns suggesting that they may also encode sRNAs (enrichment in co-IP signals at the 3' region of the gene, shown here for *tdcG* and *ygaM*).

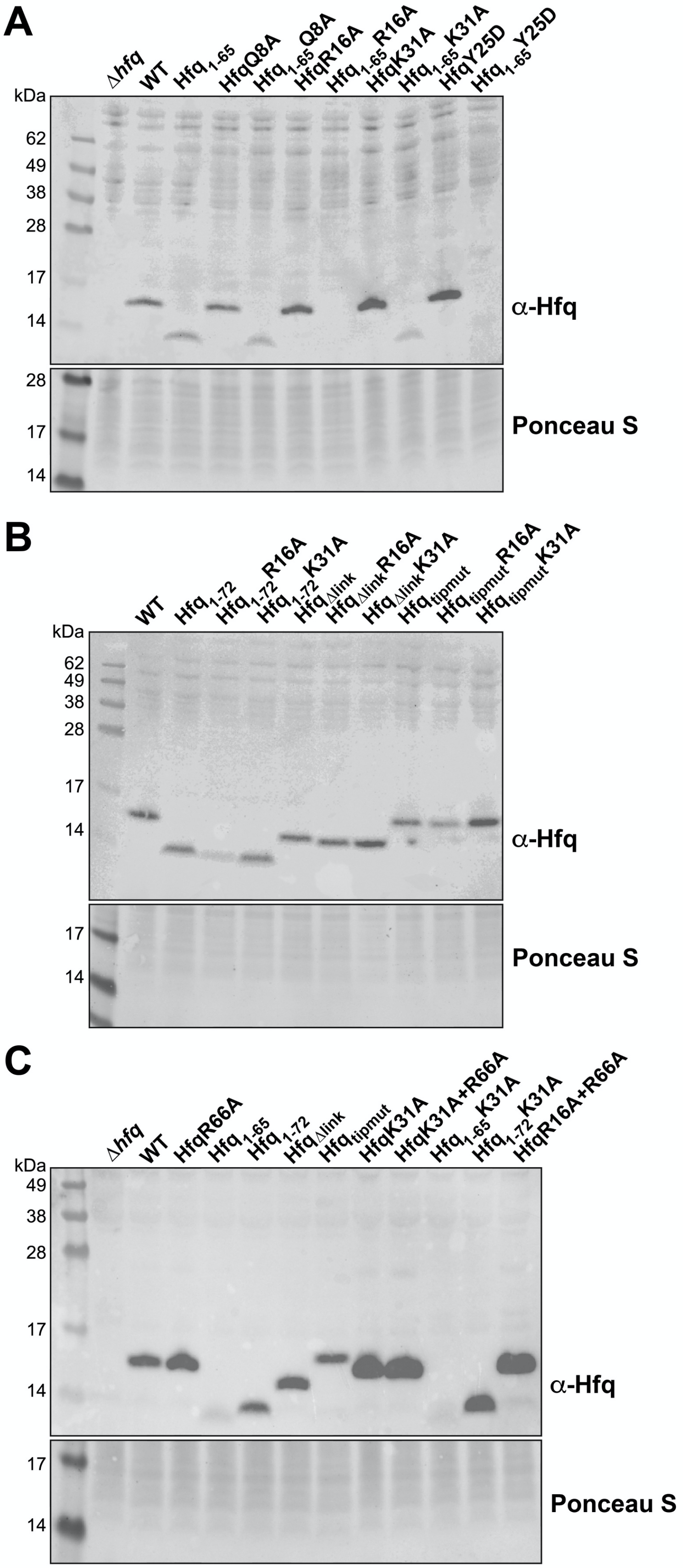
A
Δhfq
WT
Hfq1-65
HfqQ8A
Hfq1-65 Q8A
HfqR16A
Hfq1-65 R16A
HfqK31A
Hfq1-65 K31A
HfqY25D
Hfq1-65 Y25D
kDa
62
49
38
28
17
14
α-Hfq
28
17
14
Ponceau S
B
WT
Hfq1-72
Hfq1-72 R16A
Hfq1-72 K31A
HfqΔlink
HfqΔlink R16A
HfqΔlink K31A
Hfqtipmut
Hfqtipmut R16A
Hfqtipmut K31A
kDa
62
49
38
28
17
14
α-Hfq
17
14
Ponceau S
C
Δhfq
WT
HfqR66A
Hfq1-65
Hfq1-72
HfqΔlink
Hfqtipmut
HfqK31A
HfqK31A+R66A
Hfq1-65 K31A
Hfq1-72 K31A
HfqR16A+R66A
kDa
49
38
28
17
14
α-Hfq
17
14
Ponceau S

**Figure S3.** Levels of Hfq single and double mutants characterized in study.

Western blot of isogenic Hfq mutants. The strains are those used to measure *chiP-lacZ* activity in Figures 2B, 3B, 3E and 3F. (**A**) Δ*hfq* (DJS2689), WT (DJS2690), $hfq_{1\text{-}65}$ (KK01), *hfqQ8A* (DJS2691), $hfq_{1\text{-}65}$ *Q8A* (KK2414), *hfqR16A* (DJS2693), $hfq_{1\text{-}65}$ *R16A* (KK02), *hfqK31A* (DJS2695), $hfq_{1\text{-}65}$ *K31A* (KK04), *hfqY25D* (DJS2694), $hfq_{1\text{-}65}$*Y25D* (KK03), (**B**) WT (DJS2690), $hfq_{1\text{-}72}$ (KK2438), $hfq_{1\text{-}72}$*R16A* (KK2657), $hfq_{1\text{-}72}$*K31A* (KK2658), $hfq_{\Delta link}$ (KK2558) $hfq_{\Delta link}$ *R16A* (KK2654), $hfq_{\Delta link}$ *K31A* (KK2655), $hfq_{tipmut}$ (KK2570), $hfq_{tipmut}$ *R16A* (KK2651), $hfq_{tipmut}$ *K31A* (KK2652) and (**C**) Δ*hfq* (DJS2689), WT (DJS2690), *hfqR66A* (KK2725), $hfq_{1\text{-}65}$ (KK01), $hfq_{1\text{-}72}$ (KK2438), $hfq_{\Delta \text{link}}$ (KK2558), $hfq_{\text{tipmut}}$ (KK2570), *hfq*K31A (DJS2695), *hfqK31A R66A* (KK2727), $hfq_{1\text{-}65}$ *K31A* (KK04), $hfq_{1\text{-}72}$ *K31A* (DD2658), *hfqR16A R66A* (KK2726). The anti-Hfq antibody was used (upper panel) and Ponceau S as loading control (lower panel). Total protein from 1 ml of culture at $OD_{600} = 1$ was assayed. The sample preparation and antibody used are as reported previously (1). Our earlier report (2) suggests that Hfq antibody does not recognize the $Hfq_{1\text{-}65}$ monomer well, possibly due to loss of critical epitopes affecting the interpretation of the relative amounts of Hfq.

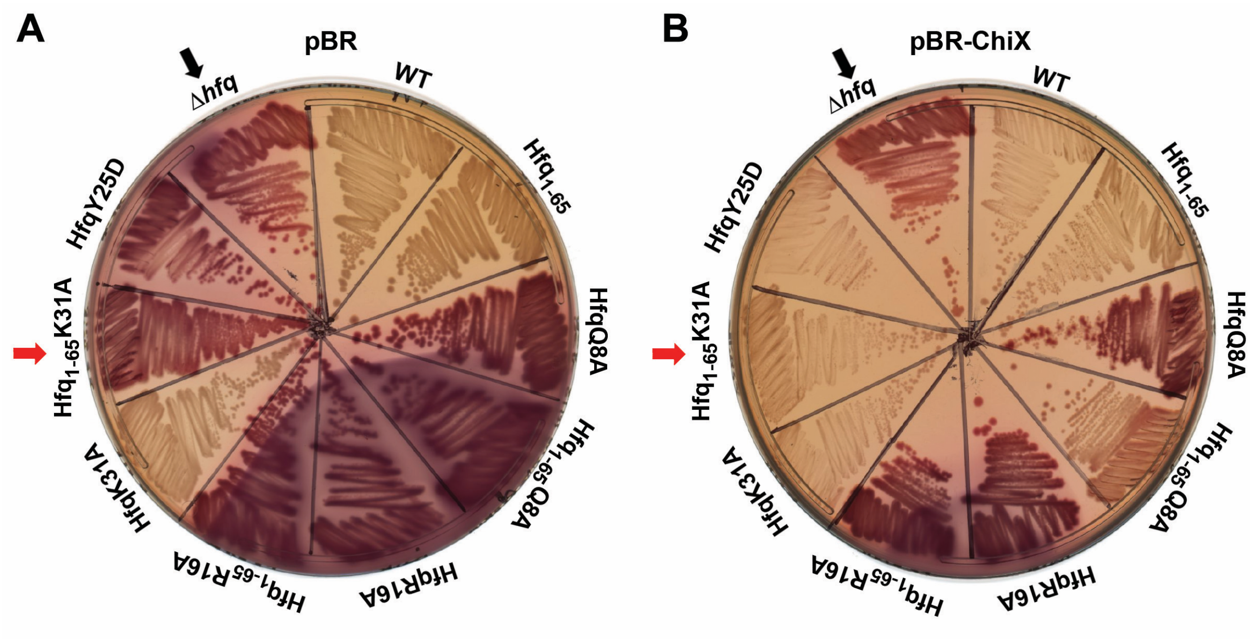


**Figure S4.** Multicopy ChiX can suppress the regulatory defect for ChiX regulation of *chiP-lacZ* in a $Hfq_{1-65}$ K31A background. Regulation of *pBAD-chiP-lacZ* [Δ*hfq* (DJS2689), *WT* (DJS2690), *hfq*$_{1-65}$ (KK01), *hfqQ8A* (DJS2691), *hfq*$_{1-65}$ *Q8A* (KK2414), *hfqR16A* (DJS2693), *hfq*$_{1-65}$ *R16A* (KK02), *hfqK31A* (DJS2695), *hfq*$_{1-65}$ *K31A* (KK04), *hfqY25D* (DJS2694)] in the presence of ChiX expressed from the chromosome with vector (pBR-plac) plasmid (**A**), or from a plasmid (**B**) (pBR-ChiX). The assay was performed on MacConkey plates with 0.0005% arabinose and 50 µg ampicillin incubated at 37˚C for 16 h. All Hfq mutants were expressed from the native *hfq* locus.

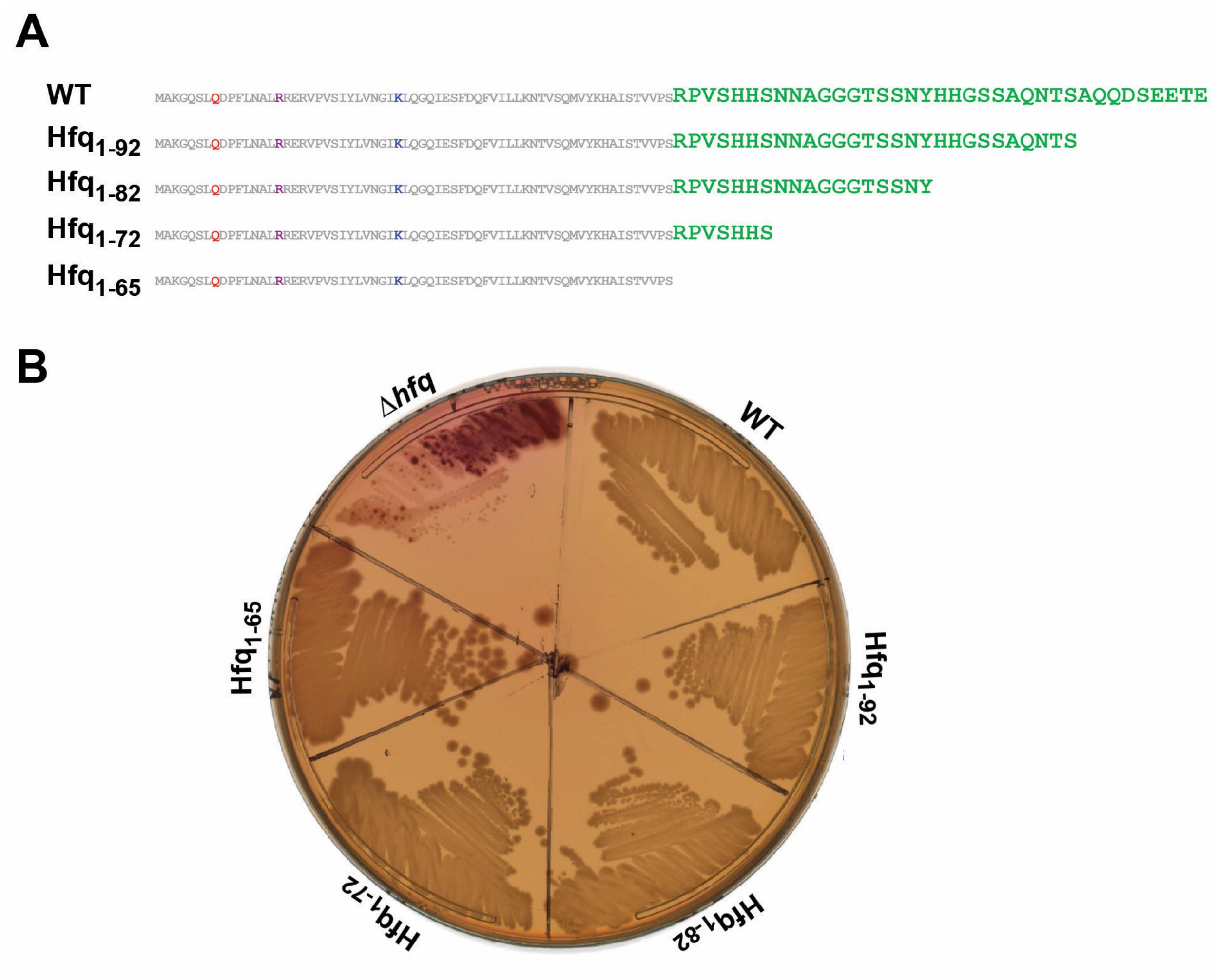


**Figure S5.** Effect of Hfq CTD truncations on ChiX-based regulation of *chiP-lacZ*. (**A**) Schematic of CTD truncations (**B**) Truncated HfqCTD mutant derivatives of PM1205 (*lacI'::P$_{BAD}$-chiP-lacZ* ), expressing ChiX from the chromosome, were constructed and the activity of P$_{BAD}$-*chiP-lacZ* measured on MacConkey lactose plates containing 0.0005% arabinose incubated at 37˚C for 16 h. Strains used: Δ*hfq* (DJS2689), *WT* (DJS2690), $hfq_{1\text{-}65}$ (KK01), $hfq_{1\text{-}72}$ (KK2438), $hfq_{1\text{-}82}$ (KK2437), $hfq_{1\text{-}92}$ (KK2436).

**Table S1.** Strains used in this study.

| Strains | Description | Reference or Source |
|---|---|---|
| AZ237 | MC4100 *hfq*+ | (3) |
| AZ238 | MC4100 *hfq*Q8A | (3) |
| AZ241 | MC4100 *hfq*K31A | (3) |
| C600 | $F^-$ *tonA21 thi-1 thr-1 leuB6 lacY1 glnV44 rfbC1 fhuA1* $\lambda^-$ | Lab collection |
| DJS2255 | PM1205 *lacI'::PBAD-ompX-lacZ Δhfq::trpAterm-kan-*$P_{BAD}$*-ccdB miniλ::tet* | (3) |
| DJS2286 | *Δhfq::cat-sacB ΔpurA::kan* | (3) |
| DJS2604 | MG1655*Δhfq::cat-sacB ΔpurA::kan* | MG1655 + P1(DJS2286) |
| DJS2609 | MG1655 $hfq^+$ | DJS2604 + P1(AZ237) |
| DJS2676 | PM1205 *lacI'::pBAD-sodB-lacZ, Δhfq:: cat-sacB::ΔpurA::kan* | (3) |
| DJS2677 | PM1205 *lacI'::*$P_{BAD}$ *-chiP-lacZ Δhfq::cat-sacB ΔpurA::kan* | (3) |
| DJS2682 | PM1205 *lacI'::pBAD-sodB-lacZ, Δhfq:: cat-sacB* | (3) |
| DJS2683 | PM1205 *lacI'::pBAD-sodB-lacZ,* $hfq^+$ | (3) |
| DJS2684 | PM1205 *lacI'::pBAD-sodB-lacZ, hfqQ8A* | (3) |
| DJS2686 | PM1205 *lacI'::pBAD-sodB-lacZ, hfqR16A* | (3) |
| DJS2687 | PM1205 *lacI'::pBAD-sodB-lacZ, hfqY25D* | (3) |
| DJS2688 | PM1205 *lacI'::pBAD-sodB-lacZ, hfqK31A* | (3) |
| DJS2689 | PM1205 *lacI'::*$P_{BAD}$*-chiP-lacZ Δhfq::cat-sacB* | (3) |
| DJS2690 | PM1205 *lacI'::*$P_{BAD}$*-chiP-lacZ* $hfq^+$(WT) | (3) |
| DJS2691 | PM1205 *lacI'::*$P_{BAD}$*-chiP-lacZ hfqQ8A* | (3) |
| DJS2693 | PM1205 *lacI'::*$P_{BAD}$*-chiP-lacZ hfqR16A* | (3) |
| DJS2694 | PM1205 *lacI'::*$P_{BAD}$*-chiP-lacZ hfqY25D* | (3) |
| DJS2695 | PM1205 *lacI'::*$P_{BAD}$*-chiP-lacZ hfqK31A* | (3) |
| DJS2814 | MG1655 *ΔlacX74. mal::lacIq ΔaraBAD* | (4) |

| | | |
|---|---|---|
| | *Δhfq::trpAterm-kan-*$P_{BAD}$*-ccdB miniλ::tet* | |
| DJS2985 | PM1205 *lacI'::*$P_{BAD}$*-chiPsodBbplacZ* | (4) |
| DJS2925 | C600 *Δhfq:: cat-sacBΔpurA::kan* | C600 + P1(DJS2286) |
| DJS3007 | PM1205 *lacI'::*$p_{BAD}$*-chiPsodB-lacZ Δhfq:: cat-sacB* | (4) |
| DJS3008 | PM1205 *lacI'::*$p_{BAD}$*-chiPsodB-lacZ, Δhfq:: cat-sacBΔpurA::kan* | (4) |
| DJS3009 | PM1205 *lacI'::*$p_{BAD}$*-chiPsodB-lacZ, hfq*$^{+}$ | (4) |
| DJS3010 | PM1205 *lacI'::*$p_{BAD}$*-chiPsodB-lacZ, hfqR16A* | (4) |
| DJS3011 | PM1205 *lacI'::*$p_{BAD}$*-chiPsodB-lacZ, hfqY25D* | (4) |
| JC1060 | *MG1655 mal::lacIQ ΔaraBAD araC+ lacI'::kan-Cp12b-mutS-lacZ* | (5) |
| JC1316 | MG1655 *lacI*$^{Q}$ *Δfur::zeo* | (6) |
| KK01 | PM1205 *lacI'::*$P_{BAD}$*-chiP-lacZ* *hfq*$_{1-65}$ | (2) |
| KK02 | PM1205 *lacI'::*$P_{BAD}$*-chiP-lacZ* *hfq*$_{1-65}$ *R16A* | DJS2677+P1(RAF1049) |
| KK03 | PM1205 *lacI'::*$P_{BAD}$*-chiP-lacZ* *hfq*$_{1-65}$ *Y25D* | DJS2677+P1(RAF1050) |
| KK04 | PM1205 *lacI'::*$P_{BAD}$*-chiP-lacZ* *hfq*$_{1-65}$ *K31A* | DJS2677 + P1(RAF1047) |
| KK05 | MG1655 *Δhfq:: cat-sacBΔpurA::kan* | MG1655 + P1(DJS2925) |
| KK2414 | PM1205 *lacI'::*$P_{BAD}$*-chiP-lacZ* *hfq*$_{1-65}$ *Q8A* | DJS2677+P1(RAF1048) |
| KK2436 | PM1205 *lacI'::*$P_{BAD}$*-chiP-lacZ, hfq*$_{1-92}$ | DJS2677+P1(RAF1002) |
| KK2437 | PM1205 *lacI'::*$P_{BAD}$*-chiP-lacZ, hfq*$_{1-82}$ | DJS2677+P1(RAF1001) |
| KK2438 | PM1205 *lacI'::*$P_{BAD}$*-chiP-lacZ, hfq*$_{1-72}$ | DJS2677+P1(RAF1000) |
| KK2440 | MG1655 *hfq*$^{+}$ | KK05 + P1(DJS2609) |
| KK2446 | C600 *hfq*$_{1-65}$ | DJS2925 + P1(RAF1042) |
| KK2448 | MG1655 *hfq*$_{1-65}$ | KK05 + P1(KK2446) |
| KK2455 | PM1205 *lacI'::kan-Cp12b-mutS-lacZ, Δhfq:: cat-sacB* | DJS2689+P1(JC1060) |
| KK2456 | PM1205 *lacI'::kan-Cp12b-mutS-lacZ, hfq*$^{+}$ | DJS2690+P1(JC1060) |
| KK2457 | PM1205 *lacI'::kan-Cp12b-mutS-lacZ, hfq*$_{1-65}$ | KK01+P1(JC1060) |
| KK2460 | PM1205 *lacI'::kan-Cp12b-mutS-lacZ, hfqQ8A* | DJS2691+P1(JC1060) |
| KK2461 | PM1205 *lacI'::kan-Cp12b-mutS-lacZ, hfq*$_{1-65}$ *Q8A* | KK2441+P1(JC1060) |
| KK2462 | PM1205 *lacI'::kan-Cp12b-mutS-lacZ, hfqR16A* | DJS2693+P1(JC1060) |

| | | |
|---|---|---|
| KK2463 | PM1205 *lacI'::kan-Cp12b-mutS-lacZ, hfq*$_{1-65}$ *R16A* | KK02+ P1(JC1060) |
| KK2464 | PM1205 *lacI'::kan-Cp12b-mutS-lacZ, hfqY25D* | DJS2694+P1(JC1060) |
| KK2466 | PM1205 *lacI'::kan-Cp12b-mutS-lacZ, hfqK31A* | DJS2695+P1(JC1060) |
| KK2467 | PM1205 *lacI'::kan-Cp12b-mutS-lacZ, hfq*$_{1-65}$ *K31A* | KK04+ P1(JC1060) |
| KK2540 | MG1655 *ΔlacX74 mal::lacIq ΔaraBAD hfq* $_{\Delta link}$ | DJS2814 + *hfq*$_{\Delta link}$ gblock, recombineering[1] |
| KK2541 | MG1655 *ΔlacX74 mal::lacIq ΔaraBAD hfq*$_{TipMut}$ | DJS2814+*TipMut*-CTD gblock, recombineering[1] |
| KK2558 | PM1205 *lacI'::P*$_{BAD}$*-chiP-lacZ hfq*$_{\Delta link}$ | DJS2677+P1(KK2540) |
| KK2570 | PM1205 *lacI'::P*$_{BAD}$*-chiP-lacZ hfq*$_{TipMut}$ | DJS2677+P1(KK2541) |
| KK2611 | PM1205 *lacI'::pBAD-sodB-lacZ,hfq*$_{1-65}$ | DJS2676+P1(KK01) |
| KK2613 | PM1205 *lacI'::pBAD-sodB-lacZ,hfq*$_{1-65}$ *Q8A* | DJS2676+P1(RAF1048) |
| KK2613r* | PM1205 *lacI'::pBAD-sodB-lacZ,hfq*$_{1-65}$ *Q8A* | Reconstructed, as for KK2613 |
| KK2615 | PM1205 *lacI'::pBAD-sodB-lacZ, hfq*$_{1-65}$ *R16A* | DJS2676+P1(RAF1049) |
| KK2619 | PM1205 *lacI'::pBAD-sodB-lacZ, hfq*$_{1-65}$ *K31A* | DJS2676+P1(RAF1047) |
| KK2622 | PM1205 *lacI'::pBAD-chiPsodB-lacZ, hfq*$_{1-65}$ | DJS3008+P1(KK01) |
| KK2623 | PM1205 *lacI'::pBAD-chiPsodB-lacZ, hfqQ8A* | DJS3008 + P1(AZ238) |
| KK2624 | PM1205 *lacI'::pBAD-chiPsodB-lacZ, hfq*$_{1-65}$ *Q8A* | DJS3008+P1(RAF1048) |
| KK2624r* | PM1205 *lacI'::pBAD-chiPsodB-lacZ, hfq*$_{1-65}$ *Q8A* | Reconstructed, as for KK2624 |
| KK2626 | PM1205 *lacI'::pBAD-chiPsodB-lacZ, hfq*$_{1-65}$ *R16A* | DJS3008+P1(RAF1049) |
| KK2629 | PM1205 *lacI'::pBAD-chiPsodB-lacZ, hfqK31A* | DJS3008+P1(AZ241) |
| KK2630 | PM1205 *lacI'::pBAD-chiPsodB-lacZ, hfq*$_{1-65}$ *K31A* | DJS3008+P1(RAF1047) |
| KK2646 | MG1655 *ΔlacX74 mal::lacIq ΔaraBAD hfq*$_{TipMut}$*R16A* | DJS2814 + *hfq*$_{TipMut}$*R16A* gblock, recombineering[1] |
| KK2647 | MG1655 *ΔlacX74 mal::lacIq ΔaraBAD hfq*$_{TipMut}$*K31A* | DJS2814 + *hfq*$_{TipMut}$*K31A* gblock recombineering[1] |
| KK2649 | MG1655 *ΔlacX74 mal::lacIq ΔaraBAD hfq*$_{\Delta link}$ R16A | DJS2814 + *hfq*$_{\Delta link}$*R16A* gblock, recombineering[1] |
| KK2650 | MG1655 *ΔlacX74 mal::lacIq ΔaraBAD hfq*$_{\Delta link}$*K31A* | DJS2814 + *hfqK31A*$_{\Delta link}$ gblock, recombineering[1] |
| KK2651 | PM1205 *lacI'::P*$_{BAD}$*-chiP-lacZ hfq*$_{TipMut}$*R16A* | DJS2677+P1(KK2646) |

| | | |
|---|---|---|
| KK2652 | PM1205 *lacI'::*$P_{BAD}$*-chiP-lacZ* $hfq_{TipMut}$*K31A* | DJS2677+P1(KK2647) |
| KK2654 | PM1205 *lacI'::*$P_{BAD}$*-chiP-lacZ* $hfq_{\Delta link}$*R16A* | DJS2677+P1(KK2649) |
| KK2655 | PM1205 *lacI'::*$P_{BAD}$*-chiP-lacZ* $hfq_{\Delta link}$*K31A* | DJS2677+P1(KK2650) |
| KK2657 | PM1205 *lacI'::*$P_{BAD}$*-chiP-lacZ,* $hfq_{1-72}$*R16A* | DJS2677+P1(RAF1053) |
| KK2658 | PM1205 *lacI'::*$P_{BAD}$*-chiP-lacZ,* $hfq_{1-72}$*K31A* | DJS2677+P1(RAF1051) |
| KK2659 | PM1205 *lacI'::pBAD-sodB-lacZ,* $hfq_{TipMut}$ | DJS2676+P1(KK2541) |
| KK2660 | PM1205 *lacI'::pBAD-sodB-lacZ,* $hfq_{\Delta link}$ | DJS2676+P1(KK2540) |
| KK2661 | PM1205 *lacI'::pBAD-sodB-lacZ,* $hfq_{TipMut}$*R16A* | DJS2676+P1(KK2646) |
| KK2662 | PM1205 *lacI'::pBAD-sodB-lacZ,* $hfq_{TipMut}$*K31A* | DJS2676+P1(KK2647) |
| KK2664 | PM1205 *lacI'::pBAD-sodB-lacZ,* $hfq_{\Delta link}$*R16A* | DJS2676+P1(KK2649) |
| KK2665 | PM1205 *lacI'::pBAD-sodB-lacZ,* $hfq_{\Delta link}$*K31A* | DJS2676+P1(KK2650) |
| KK2667 | PM1205 *lacI'::pBAD-sodB-lacZ,* $Hfq_{1-72}$*R16A* | DJS2676+P1(RAF1053) |
| KK2668 | PM1205 *lacI'::pBAD-sodB-lacZ* $hfq_{1-72}$*K31A* | DJS2676+P1(RAF1051) |
| KK2674 | MG1655 *ΔlacX74 mal::lacIq ΔaraBAD* $hfq_{1-72}$*R66AK31A* | DJS2814+$hfq_{1-72}$R66AK31A gblock, recombineering[1] |
| KK2675 | MG1655 *ΔlacX74 mal::lacIq ΔaraBAD* $hfq_{1-72}$*P67AK31A* | DJS2814+$hfq_{1-72}$P67AK31A gblock, recombineering[1] |
| KK2676 | MG1655 *ΔlacX74 mal::lacIq ΔaraBAD* $hfq_{1-72}$*H70AK31A* | DJS2814+$hfq_{1-72}$H70AK31A gblock, recombineering[1] |
| KK2677 | MG1655 *ΔlacX74 mal::lacIq ΔaraBAD* $hfq_{1-72}$*H71AK31A* | DJS2814+$hfq_{1-72}$H71AK31A gblock, recombineering[1] |
| KK2678 | MG1655 *ΔlacX74 mal::lacIq ΔaraBAD* $hfq_{1-72}$*S72AK31A* | DJS2814+$hfq_{1-72}$S72AK31A gblock, recombineering[1] |
| KK2680 | PM1205 *lacI'::*$P_{BAD}$*-chiP-lacZ,* $hfq_{1-72}$*R66AK31A* | DJS2677+P1(KK2674) |
| KK2681 | PM1205 *lacI'::*$P_{BAD}$*-chiP-lacZ,* $hfq_{1-72}$*P67AK31A* | DJS2677+P1(KK2675) |
| KK2682 | PM1205 *lacI'::*$P_{BAD}$*-chiP-lacZ,* $hfq_{1-72}$*H70AK31A* | DJS2677+P1(KK2676) |
| KK2683 | PM1205 *lacI'::*$P_{BAD}$*-chiP-lacZ,* $hfq_{1-72}$*H71AK31A* | DJS2677+P1(KK2677) |
| KK2684 | PM1205 *lacI'::*$P_{BAD}$*-chiP-lacZ,* $hfq_{1-72}$*S72AK31A* | DJS2677+P1(KK2678) |

| | | |
|---|---|---|
| KK2685 | MG1655 *ΔlacX74 mal::lacIq ΔaraBAD* $hfq_{1\text{-}72}$*V68AK31A* | DJS2814+$hfq_{1\text{-}72}$V68AK31A gblock, recombineering[1] |
| KK2686 | MG1655 *ΔlacX74 mal::lacIq ΔaraBAD* $hfq_{1\text{-}72}$*S69AK31A* | DJS2814+$hfq_{1\text{-}72}$S69AK31A gblock, recombineering[1] |
| KK2687 | PM1205 *lacI'::*$P_{BAD}$*-chiP-lacZ,* $hfq_{1\text{-}72}$*V68AK31A* | DJS2677+P1(KK2685) |
| KK2688 | PM1205 *lacI'::*$P_{BAD}$*-chiP-lacZ,* $hfq_{1\text{-}72}$*S69AK31A* | DJS2677+P1(KK2686) |
| KK2689 | PM1205 *lacI'::kan-Cp12b-mutS-lacZ,* $hfq_{1\text{-}72}$ | KK2438+ P1(JC1060) |
| KK2691 | PM1205 *lacI'::kan-Cp12b-mutS-lacZ,* $hfq_{1\text{-}72}$*R16A* | KK2657+P1(JC1060) |
| KK2692 | PM1205 *lacI'::kan-Cp12b-mutS-lacZ,* $hfq_{1\text{-}72}$*K31A* | KK2658+ P1(JC1060) |
| KK2693 | PM1205 *lacI'::pBAD-sodB-lacZ, Δfur::zeo,* $hfq^{+}$*/WT* | DJS2683+ P1(JC1316) |
| KK2694 | PM1205 *lacI'::pBAD-sodB-lacZ Δfur::zeo,* $hfq_{1\text{-}65}$ | KK2611+ P1(JC1316) |
| KK2695 | PM1205 *lacI'::pBAD-sodB-lacZ Δfur::zeo, hfqQ8A* | DJS2684+ P1(JC1316) |
| KK2696 | PM1205 *lacI'::pBAD-sodB-lacZ Δfur::zeo, hfqR16A* | DJS2686+ P1(JC1316) |
| KK2697 | PM1205 *lacI'::pBAD-sodB-lacZ Δfur::zeo,* $hfq_{\Delta link}$*R16A* | KK2664+ P1(JC1316) |
| KK2698 | PM1205 *lacI'::pBAD-sodB-lacZ Δfur::zeo,* $hfq_{tipmut}$*R16A* | KK2661+ P1(JC1316) |
| KK2699 | PM1205 *lacI'::pBAD-sodB-lacZ, Δfur::zeo,* $hfq_{\Delta link}$ | KK2660+P1(JC1316) |
| KK2700 | PM1205 *lacI'::pBAD-sodB-lacZ, Δfur::zeo,* $hfq_{tipmut}$ | KK2659+P1(JC1316) |
| KK2701 | PM1205 *lacI'::pBAD-sodB-lacZ Δfur::zeo, hfqK31A* | DJS2688+ P1(JC1316) |
| KK2702 | PM1205 *lacI'::pBAD-sodB-lacZ Δfur::zeo,* $hfq_{1\text{-}65}$ *K31A* | KK2619+ P1(JC1316) |
| KK2704 | PM1205 *lacI'::pBAD-sodB-lacZ Δfur::zeo,* $hfq_{1\text{-}72}$*R16A* | KK2667+P1(JC1316) |
| KK2705 | PM1205 *lacI'::pBAD-sodB-lacZ Δfur::zeo,* $hfq_{1\text{-}72}$*K31A* | KK2668+P1(JC1316) |
| KK2706 | PM1205 *lacI'::pBAD-sodB-lacZ Δfur::zeo, Δhfq:: cat-sacB* | DJS2682+P1(JC1316) |
| KK2707 | PM1205 *lacI'::pBAD-sodB-lacZ Δfur::zeo,* $hfq_{1\text{-}65}$ *R16A* | KK2615+ P1(JC1316) |
| KK2708 | PM1205 *lacI'::pBAD-sodB-lacZ,* $hfq_{1\text{-}72}$ | DJS2676+P1(RAF1000) |
| KK2714 | PM1205 *lacI'::pBAD-sodB-lacZ Δfur::zeo,* $hfq_{tipmut}$*K31A* | KK2662+ P1(JC1316) |

| | | |
|---|---|---|
| KK2715 | PM1205 *lacI'::pBAD-sodB-lacZ Δfur::zeo,hfq*$_{\Delta link}$*K31A* | KK2665+ P1(JC1316) |
| KK2717 | PM1205 *lacI'::pBAD-sodB-lacZ Δfur::zeo, hfq*$_{1\text{-}65}$ *Q8A* | KK2613+ P1(JC1316) |
| KK2717r* | PM1205 *lacI'::pBAD-sodB-lacZ Δfur::zeo, hfq*$_{1\text{-}65}$ *Q8A* | Reconstructed, as for KK2717 |
| KK2718 | PM1205 *lacI'::pBAD-sodB-lacZ Δfur::zeo, hfq*$_{1\text{-}72}$ | KK2708+P1(JC1316) |
| KK2719 | PM1205 *lacI'::pBAD-sodB-lacZ Δfur::zeo, hfqY25D* | DJS2687+ P1(JC1316) |
| KK2722 | MG1655 *ΔlacX74 mal::lacIq ΔaraBAD hfqR66A* | DJS2814 + R66A gblock, recombineering[1] |
| KK2723 | MG1655 *ΔlacX74mal::lacIq ΔaraBAD hfqR66AR16A* | DJS2814 + R66AR16A gblock, recombineering[1] |
| KK2724 | MG1655 *ΔlacX74 mal::lacIq ΔaraBAD hfqR66AK31A* | DJS2814 + R66AK31A gblock, recombineering[1] |
| KK2725 | PM1205 *lacI'::pBAD-chiP-lacZ, hfq*R66A | DJS2677+P1(KK2722) |
| KK2726 | PM1205 *lacI'::pBAD-chiP-lacZ, hfq*R66AR16A | DJS2677+P1(KK2723) |
| KK2727 | PM1205 *lacI'::pBAD-chiP-lacZ, hfq*R66AK31A | DJS2677+P1(KK2724) |
| KK2728 | PM1205 *lacI'::pBAD-sodB-lacZ , hfqR66A* | DJS2676+P1(KK2722) |
| KK2729 | PM1205 *lacI'::pBAD-sodB-lacZ, hfqR66AR16A* | DJS2676+P1(KK2723) |
| KK2730 | PM1205 *lacI'::pBAD-sodB-lacZ, hfqR66AK31A* | DJS2676+P1(KK2724) |
| KK2731 | PM1205 *lacI'::pBAD-sodB-lacZ Δfur::zeo, hfqR66A* | KK2728+ P1(JC1316) |
| KK2732 | PM1205 *lacI'::pBAD-sodB-lacZ Δfur::zeo, hfqR66AR16A* | KK2729+ P1(JC1316) |
| KK2732r* | PM1205 *lacI'::pBAD-sodB-lacZ Δfur::zeo, hfqR66AR16A* | Reconstructed as for KK2732 |
| KK2733 | PM1205 *lacI'::pBAD-sodB-lacZ Δfur::zeo, hfqR66AK31A* | KK2730+ P1(JC1316) |
| KK2733r* | PM1205 *lacI'::pBAD-sodB-lacZ Δfur::zeo, hfqR66AK31A* | Reconstructed as for KK2733 |
| PM1205 | *mal::lacI*$^q$, *araC*$^+$, $P_{BAD}$*-cat-sacB-lacZ*, mini λ tet$^R$ | (7) |
| RAF1000 | PM1205 *lacI'::*$P_{BAD}$*-ompX-lacZ hfq*$_{1\text{-}72}$ | Hfq72 PCR fragment amplified from pNRD414 with Aztunc1000 forward and Hfq72 reverse primers + DJS2255, recombineering. [1] |

| | | |
|---|---|---|
| RAF1001 | PM1205 *lacI'::$P_{BAD}$-ompX-lacZ hfq*$_{1\text{-}82}$ | Hfq82 PCR fragment amplified from pNRD414 with Aztunc1000 forward and Hfq82 reverse primers + DJS2255, recombineering. [1] |
| RAF1002 | PM1205 *lacI'::$P_{BAD}$-ompX-lacZ hfq*$_{1\text{-}92}$ | Hfq 92 PCR fragment amplified from pNRD414 with Aztunc1000 forward and Hfq92 reverse primers + DJS2255, recombineering. [1] |
| RAF1042 | PM1205 *lacI'::$P_{BAD}$-ompX-lacZ hfq*$_{1\text{-}65}$ | Santiago-Frangos et al 2016 |
| RAF1047 | PM1205 *lacI'*::$P_{BAD}$-*ompX-lacZ hfq*$_{1\text{-}65}$ *K31A* | Hfq 65K31A PCR fragment amplified from DJS2695 with Aztunc1000 forward and Hfq65 reverse primers + DJS2255, recombineering. [1] |
| RAF1048 | PM1205 *lacI'::$P_{BAD}$-ompX-lacZ hfq*$_{1\text{-}65}$ *Q8A* | Hfq 65Q8A PCR fragment amplified from DJS2691 with Aztunc1000 forward and Hfq65 reverse primers + DJS2255, recombineering. [1] |
| RAF1049 | PM1205 *lacI'::$P_{BAD}$-ompX-lacZ hfq*$_{1\text{-}65}$ *R16A* | Hfq 65R16A PCR fragment amplified from DJS2693 with Aztunc1000 forward and Hfq65 reverse primers + DJS2255, recombineering. [1] |
| RAF1050 | PM1205 *lacI'::$P_{BAD}$-ompX-lacZ hfq*$_{1\text{-}65}$ *Y25D* | Hfq 65Y25D PCR fragment amplified from DJS2694 with Aztunc1000 forward and Hfq65 reverse primers + DJS2255 recombineering. [1] |
| RAF1051 | PM1205 *lacI'::$P_{BAD}$-ompX-lacZ hfq*$_{1\text{-}72}$*K31A* | Hfq72K31APCR fragment amplified from DJS2695 with Aztunc1000 forward and Hfq72 reverse primers + DJS2255 recombineering. [1] |
| RAF1053 | PM1205 *lacI'::$P_{BAD}$-ompX-lacZ hfq*$_{1\text{-}72}$*R16A* | Hfq 72R16A PCR fragment amplified from DJS2693 with Aztunc1000 forward and Hfq72reverse primers + DJS2255, recombineering. [1] |

[1]Using lambda red recombineering, as described in Materials and Methods, DNA fragments (gblocks or PCR fragments) with flanking homology to *hfq* were introduced into the bacterial chromosome at the native *hfq* locus.  The recipient strains carry counterelectable markers in the *hfq* locus.

**Table S2.** Oligonucleotides used in this study.

| **Name** | **Oligonucleotide Sequence** |
|---|---|
| **gBlocks** | **Sequence (5' to 3')** |
| *hfqTipMut* | AAGGTTCAAAGTACAAATAAGCATATAAGGAAAAGAGAGAATGG CTAAGGGGCAATCTTTACAAGATCCGTTCCTGAACGCACTGCGTCG GGAACGTGTTCCAGTTTCTATTTATTTGGTGAATGGTATTAAGCTG CAAGGGCAAATCGAGTCTTTTGATCAGTTCGTGATCCTGTTGAAAA ACACGGTCAGCCAGATGGTTTACAAGCACGCGATTTCTACTGTTGT CCCGTCTCGCCCGGTTTCTCATCACAGTAACAACGCCGGTGGCGGT ACCAGCAGTAACTACCATCATGGTAGCAGCGCGCAGAATACTTCC GCGCAACAGCGTAGCAACAAAACCAACTAAGGTTTCGGGCTGTTT TTTTACACGGGGAGCCAGCGATCCT |
| $hfq_{\Delta link}$ | AAGGTTCAAAGTACAAATAAGCATATAAGGAAAAGAGAGAATGG CTAAGGGGCAATCTTTACAAGATCCGTTCCTGAACGCACTGCGTCG GGAACGTGTTCCAGTTTCTATTTATTTGGTGAATGGTATTAAGCTG CAAGGGCAAATCGAGTCTTTTGATCAGTTCGTGATCCTGTTGAAAA ACACGGTCAGCCAGATGGTTTACAAGCACGCGATTTCTACTGTTGT CCCGTCTCGCCCGGTTTCTCATCACAGTAACAACGCCGGTGGCGGT ACCAGCAGTAACTACCATCATGACAGCGAAGAAACCGAATAAGGT TTCGGGCTGTTTTTTACACGGGGAGCCAGCGATCCT |
| $hfq_{1\text{-}72}R66AK31A$ | AAGGTTCAAAGTACAAATAAGCATATAAGGAAAAGAGAGAatgGCT AAGGGGCAATCTTTACAAGATCCGTTCCTGAACGCACTGCGTCGG GAACGTGTTCCAGTTTCTATTTATTTGGTGAATGGTATTGCGCTGC AAGGGCAAATCGAGTCTTTTGATCAGTTCGTGATCCTGTTGAAAAA CACGGTCAGCCAGATGGTTTACAAGCACGCGATTTCTACTGTTGTC CCGTCTGCCCCGGTTTCTCATCACAGTtaaGGTTTCGGGCTGTTTTT TACACGGGGAGCCAGCGATCCT |
| $hfq_{1\text{-}72}P67AK31A$ | AAGGTTCAAAGTACAAATAAGCATATAAGGAAAAGAGAGAATGG CTAAGGGGCAATCTTTACAAGATCCGTTCCTGAACGCACTGCGTCG GGAACGTGTTCCAGTTTCTATTTATTTGGTGAATGGTATTGCGCTG CAAGGGCAAATCGAGTCTTTTGATCAGTTCGTGATCCTGTTGAAAA ACACGGTCAGCCAGATGGTTTACAAGCACGCGATTTCTACTGTTGT CCCGTCTCGCGCGGTTTCTCATCACAGTTAAGGTTTCGGGCTGTTT TTTTACACGGGGAGCCAGCGATCCT |
| $hfq_{1\text{-}72}V68AK31A$ | AAGGTTCAAAGTACAAATAAGCATATAAGGAAAAGAGAGAATGG CTAAGGGGCAATCTTTACAAGATCCGTTCCTGAACGCACTGCGTCG GGAACGTGTTCCAGTTTCTATTTATTTGGTGAATGGTATTGCGCTG CAAGGGCAAATCGAGTCTTTTGATCAGTTCGTGATCCTGTTGAAAA ACACGGTCAGCCAGATGGTTTACAAGCACGCGATTTCTACTGTTGT CCCGTCTCGCCCGGCTTCTCATCACAGTTAAGGTTTCGGGCTGTTT TTTTACACGGGGAGCCAGCGATCCT |

*hfq*$_{1\text{-}72}$*S69AK31A*

AAGGTTCAAAGTACAAATAAGCATATAAGGAAAAGAGAGAATGG
CTAAGGGGCAATCTTTACAAGATCCGTTCCTGAACGCACTGCGTCG
GGAACGTGTTCCAGTTTCTATTTATTTGGTGAATGGTATTGCGCTG
CAAGGGCAAATCGAGTCTTTTGATCAGTTCGTGATCCTGTTGAAAA
ACACGGTCAGCCAGATGGTTTACAAGCACGCGATTTCTACTGTTGT
CCCGTCTCGCCCGGTTGCTCATCACAGTTAAGGTTTCGGGCTGTTT
TTTTACACGGGGAGCCAGCGATCCT

*hfq*$_{1\text{-}72}$*H70AK31A*

AAGGTTCAAAGTACAAATAAGCATATAAGGAAAAGAGAGAATGG
CTAAGGGGCAATCTTTACAAGATCCGTTCCTGAACGCACTGCGTCG
GGAACGTGTTCCAGTTTCTATTTATTTGGTGAATGGTATTGCGCTG
CAAGGGCAAATCGAGTCTTTTGATCAGTTCGTGATCCTGTTGAAAA
ACACGGTCAGCCAGATGGTTTACAAGCACGCGATTTCTACTGTTGT
CCCGTCTCGCCCGGTTTCTGCTCACAGTTAAGGTTTCGGGCTGTTTT
TTTACACGGGGAGCCAGCGATCCT

*hfq*$_{1\text{-}72}$*H71AK31A*

AAGGTTCAAAGTACAAATAAGCATATAAGGAAAAGAGAGAATGG
CTAAGGGGCAATCTTTACAAGATCCGTTCCTGAACGCACTGCGTCG
GGAACGTGTTCCAGTTTCTATTTATTTGGTGAATGGTATTGCGCTG
CAAGGGCAAATCGAGTCTTTTGATCAGTTCGTGATCCTGTTGAAAA
ACACGGTCAGCCAGATGGTTTACAAGCACGCGATTTCTACTGTTGT
CCCGTCTCGCCCGGTTTCTCATGCCAGTTAAGGTTTCGGGCTGTTTT
TTTACACGGGGAGCCAGCGATCCT

*hfq*$_{1\text{-}72}$*S72AK31A*

AAGGTTCAAAGTACAAATAAGCATATAAGGAAAAGAGAGAATGG
CTAAGGGGCAATCTTTACAAGATCCGTTCCTGAACGCACTGCGTCG
GGAACGTGTTCCAGTTTCTATTTATTTGGTGAATGGTATTGCGCTG
CAAGGGCAAATCGAGTCTTTTGATCAGTTCGTGATCCTGTTGAAAA
ACACGGTCAGCCAGATGGTTTACAAGCACGCGATTTCTACTGTTGT
CCCGTCTCGCCCGGTTTCTCATCACGCTTAAGGTTTCGGGCTGTTTT
TTTACACGGGGAGCCAGCGATCCT

*hfqR66A*

AAGGTTCAAAGTACAAATAAGCATATAAGGAAAAGAGAGAAtgGC
TAAGGGGCAATCTTTACAAGATCCGTTCCTGAACGCACTGCGTCG
GGAACGTGTTCCAGTTTCTATTTATTTGGTGAATGGTATTAAGCTG
CAAGGGCAAATCGAGTCTTTTGATCAGTTCGTGATCCTGTTGAAAA
ACACGGTCAGCCAGATGGTTTACAAGCACGCGATTTCTACTGTTGT
CCCGTCTGCCCCGGTTTCTCATCACAGTAACAACGCCGGTGGCGGT
ACCAGCAGTAACTACCATCATGGTAGCAGCGCGCAGAATACTTCC
GCGCAACAGGACAGCGAAGAAACCGAAtaaGGTTTCGGGCTGTTTT
TTTACACGGGGAGCCAGCGATCCT

*hfqR66AR16A*

AAGGTTCAAAGTACAAATAAGCATATAAGGAAAAGAGAGAAtgGC
TAAGGGGCAATCTTTACAAGATCCGTTCCTGAACGCACTGGCTCG
GGAACGTGTTCCAGTTTCTATTTATTTGGTGAATGGTATTAAGCTG
CAAGGGCAAATCGAGTCTTTTGATCAGTTCGTGATCCTGTTGAAAA
ACACGGTCAGCCAGATGGTTTACAAGCACGCGATTTCTACTGTTGT
CCCGTCTGCCCCGGTTTCTCATCACAGTAACAACGCCGGTGGCGGT
ACCAGCAGTAACTACCATCATGGTAGCAGCGCGCAGAATACTTCC

GCGCAACAGGACAGCGAAGAAACCGAAtaaGGTTTCGGGCTGTTTT
TTTACACGGGGAGCCAGCGATCCT

*hfqR66AK31A*

AAGGTTCAAAGTACAAATAAGCATATAAGGAAAAGAGAGAAtgGC
TAAGGGGCAATCTTTACAAGATCCGTTCCTGAACGCACTGCGTCG
GGAACGTGTTCCAGTTTCTATTTATTTGGTGAATGGTATTGCGCTG
CAAGGGCAAATCGAGTCTTTTGATCAGTTCGTGATCCTGTTGAAAA
ACACGGTCAGCCAGATGGTTTACAAGCACGCGATTTCTACTGTTGT
CCCGTCTGCCCCGGTTTCTCATCACAGTAACAACGCCGGTGGCGGT
ACCAGCAGTAACTACCATCATGGTAGCAGCGCGCAGAATACTTCC
GCGCAACAGGACAGCGAAGAAACCGAAtaaGGTTTCGGGCTGTTTT
TTTACACGGGGAGCCAGCGATCCT

*hfqQ8ATipMut*

AAGGTTCAAAGTACAAATAAGCATATAAGGAAAAGAGAGAATGG
CTAAGGGGCAATCTTTAGCAGATCCGTTCCTGAACGCACTGCGTCG
GGAACGTGTTCCAGTTTCTATTTATTTGGTGAATGGTATTAAGCTG
CAAGGGCAAATCGAGTCTTTTGATCAGTTCGTGATCCTGTTGAAAA
ACACGGTCAGCCAGATGGTTTACAAGCACGCGATTTCTACTGTTGT
CCCGTCTCGCCCGGTTTCTCATCACAGTAACAACGCCGGTGGCGGT
ACCAGCAGTAACTACCATCATGGTAGCAGCGCGCAGAATACTTCC
GCGCAACAGCGTAGCAACAAAACCAACTAAGGTTTCGGGCTGTTT
TTTTACACGGGGAGCCAGCGATCCT

*hfqR16ATipMut*

AAGGTTCAAAGTACAAATAAGCATATAAGGAAAAGAGAGAATGG
CTAAGGGGCAATCTTTACAAGATCCGTTCCTGAACGCACTGGCTCG
GGAACGTGTTCCAGTTTCTATTTATTTGGTGAATGGTATTAAGCTG
CAAGGGCAAATCGAGTCTTTTGATCAGTTCGTGATCCTGTTGAAAA
ACACGGTCAGCCAGATGGTTTACAAGCACGCGATTTCTACTGTTGT
CCCGTCTCGCCCGGTTTCTCATCACAGTAACAACGCCGGTGGCGGT
ACCAGCAGTAACTACCATCATGGTAGCAGCGCGCAGAATACTTCC
GCGCAACAGCGTAGCAACAAAACCAACTAAGGTTTCGGGCTGTTT
TTTTACACGGGGAGCCAGCGATCCT

*hfqK31ATipMut*

AAGGTTCAAAGTACAAATAAGCATATAAGGAAAAGAGAGAATGG
CTAAGGGGCAATCTTTACAAGATCCGTTCCTGAACGCACTGCGTCG
GGAACGTGTTCCAGTTTCTATTTATTTGGTGAATGGTATTGCGCTG
CAAGGGCAAATCGAGTCTTTTGATCAGTTCGTGATCCTGTTGAAAA
ACACGGTCAGCCAGATGGTTTACAAGCACGCGATTTCTACTGTTGT
CCCGTCTCGCCCGGTTTCTCATCACAGTAACAACGCCGGTGGCGGT
ACCAGCAGTAACTACCATCATGGTAGCAGCGCGCAGAATACTTCC
GCGCAACAGCGTAGCAACAAAACCAACTAAGGTTTCGGGCTGTTT
TTTTACACGGGGAGCCAGCGATCCT

*$hfq_{\Delta link}Q8A$*

AAGGTTCAAAGTACAAATAAGCATATAAGGAAAAGAGAGAATGG
CTAAGGGGCAATCTTTAGCAGATCCGTTCCTGAACGCACTGCGTCG
GGAACGTGTTCCAGTTTCTATTTATTTGGTGAATGGTATTAAGCTG
CAAGGGCAAATCGAGTCTTTTGATCAGTTCGTGATCCTGTTGAAAA
ACACGGTCAGCCAGATGGTTTACAAGCACGCGATTTCTACTGTTGT
CCCGTCTCGCCCGGTTTCTCATCACAGTAACAACGCCGGTGGCGGT

ACCAGCAGTAACTACCATCATGACAGCGAAGAAACCGAATAAGGT
TTCGGGCTGTTTTTTACACGGGGAGCCAGCGATCCT

*hfq*$_{\Delta link}$*R16A*

AAGGTTCAAAGTACAAATAAGCATATAAGGAAAAGAGAGAATGG
CTAAGGGGCAATCTTTACAAGATCCGTTCCTGAACGCACTGGCTCG
GGAACGTGTTCCAGTTTCTATTTATTTGGTGAATGGTATTAAGCTG
CAAGGGCAAATCGAGTCTTTTGATCAGTTCGTGATCCTGTTGAAAA
ACACGGTCAGCCAGATGGTTTACAAGCACGCGATTTCTACTGTTGT
CCCGTCTCGCCCGGTTTCTCATCACAGTAACAACGCCGGTGGCGGT
ACCAGCAGTAACTACCATCATGACAGCGAAGAAACCGAATAAGGT
TTCGGGCTGTTTTTTACACGGGGAGCCAGCGATCCT

*hfq*$_{\Delta link}$*K31A*

AAGGTTCAAAGTACAAATAAGCATATAAGGAAAAGAGAGAATGG
CTAAGGGGCAATCTTTACAAGATCCGTTCCTGAACGCACTGCGTCG
GGAACGTGTTCCAGTTTCTATTTATTTGGTGAATGGTATTGCGCTG
CAAGGGCAAATCGAGTCTTTTGATCAGTTCGTGATCCTGTTGAAAA
ACACGGTCAGCCAGATGGTTTACAAGCACGCGATTTCTACTGTTGT
CCCGTCTCGCCCGGTTTCTCATCACAGTAACAACGCCGGTGGCGGT
ACCAGCAGTAACTACCATCATGACAGCGAAGAAACCGAATAAGGT
TTCGGGCTGTTTTTTACACGGGGAGCCAGCGATCCT

*hfq*$_{1\text{-}65}$

AAGGTTCAAAGTACAAATAAGCATATAAGGAAAAGAGAGAATGG
CTAAGGGGCAATCTTTACAAGATCCGTTCCTGAACGCACTGCGTCG
GGAACGTGTTCCAGTTTCTATTTATTTGGTGAATGGTATTAAGCTG
CAAGGGCAAATCGAGTCTTTTGATCAGTTCGTGATCCTGTTGAAAA
ACACGGTCAGCCAGATGGTTTACAAGCACGCGATTTCTACTGTTGT
CCCGTCTTAAGGTTTCGGGCTGTTTTTTACACGGGGAGCCAGCGA
TCCT

*hfq102*

AAGGTTCAAAGTACAAATAAGCATATAAGGAAAAGAGAGAATGG
CTAAGGGGCAATCTTTACAAGATCCGTTCCTGAACGCACTGCGTCG
GGAACGTGTTCCAGTTTCTATTTATTTGGTGAATGGTATTAAGCTG
CAAGGGCAAATCGAGTCTTTTGATCAGTTCGTGATCCTGTTGAAAA
ACACGGTCAGCCAGATGGTTTACAAGCACGCGATTTCTACTGTTGT
CCCGTCTCGCCCGGTTTCTCATCACAGTAACAACGCCGGTGGCGGT
ACCAGCAGTAACTACCATCATGGTAGCAGCGCGCAGAATACTTCC
GCGCAACAGGACAGCGAAGAAACCGAATAAGGTTTCGGGCTGTTT
TTTTACACGGGGAGCCAGCGATCCT

| **Primers for strain construction** | **Sequences (5’ to 3’)** |
|---|---|
| AZ1000TrunF | AAGGTTCAAAGTACAAATAAGCATATAAGGAAAAGAGAGAATGGCTAAGGGGCAATCTTT |
| Hfq Trunc 72R | AGGATCGCTGGCTCCCCGTGTAAAAAACAGCCCGAAACCTTAACTGTGATGAGAAACCGGGC |
| Hfq Trunc 82R | AGGATCGCTGGCTCCCCGTGTAAAAAACAGCCCGAAACCTTAGTTACTGCTGGTACCGCCAC |

| | |
|---|---|
| Hfq Trunc 92R | AGGATCGCTGGCTCCCCGTGTAAAAAAACAGCCCGAAACCTTAAGT ATTCTGCGCGCTGCTAC |
| **Primers used for cDNA library preparation (RNA-Seq), no. of nucleotides** | **3' Barcode adapter sequences** |
| AZ1331, BC1, 30 | 5'P-AACATTATTAGATCGGAAGAGCGTCGTGTA-3'SpC |
| AZ1332, BC2, 30 | 5'P-AAAGTGTTGAGATCGGAAGAGCGTCGTGTA-3'SpC |
| AZ1333, BC3, 30 | 5'P-AAGAATTATAGATCGGAAGAGCGTCGTGTA-3'SpC |
| AZ1334, BC4, 30 | 5'P-AATATGGACAGATCGGAAGAGCGTCGTGTA-3'SpC |
| AZ1335, BC5, 30 | 5'P-AATCACTTGAGATCGGAAGAGCGTCGTGTA-3'SpC |
| AZ1336, BC6, 30 | 5'P-ACCAAGTCGAGATCGGAAGAGCGTCGTGTA-3'SpC |
| AZ1337, BC7, 30 | 5'P-ACAACTCGCAGATCGGAAGAGCGTCGTGTA-3'SpC |
| AZ1338, BC8, 30 | 5'P-ACCCGTCTTAGATCGGAAGAGCGTCGTGTA-3'SpC |
| AZ1339, BC9, 30 | 5'P-ACCCTACAGAGATCGGAAGAGCGTCGTGTA-3'SpC |
| AZ1340, BC10, 30 | 5'P-ACCCTCGGCAGATCGGAAGAGCGTCGTGTA-3'SpC |
| AZ1341, BC11, 30 | 5'P-ACCGGTACCAGATCGGAAGAGCGTCGTGTA-3'SpC |
| AZ1342, BC12, 30 | 5'P-ACGGAGGGCAGATCGGAAGAGCGTCGTGTA-3'SpC |
| AZ1343, 3Tr3, 22 | 5'P-AGA TCG GAA GAG CAC ACG TCT G-3'SpC |
| AZ1344, AR2, 19 | 5'P-TACACGACGCTCTTCCGAT |
| **Primers used for northern analysis** | |
| AZ1200, ChiX | GCTATTGGCCCGTCAAAGAG |
| AZ1371, MgrR | GCGGTGAATGCTTGCATGGATAGA |
| PA006, CyaR | GGGAGATTACACAGGCTAAGGAGGTGGTTCCTGGTACAGC |
| AK280, GlnZ | ATGGGCTACAGATAGCTGACAAACTTCACG |

| | |
|---|---|
| AZ1318, MicF | GCGAGGCATCCGGTTGAAATAGGGGTAAACAGACATTCAG |
| AZ1455, OmrB | CATCTGCGCAGGCTGGTGTAATTCATGTGCTCAAC |
| AZ1324, RyhB | ACTGGAAGCAATGTGAGCAATGTCGTGCT |
| PA027, 5S | CGGCGCTACGGCGTTTCACTTCTG |

**Table S3.** RNA-Seq: Levels of all RNA species in total RNA and co-IP samples for WT and $Hfq_{1-65}$. Gene names in column A and "type" (CDS, sRNA, etc.) are as determined by genome version used for the analysis. For a small number of the genes of interest discussed here, recent studies and the profile of the browser images for Hfq Co-IP data suggest the existence of unannotated sRNAs. In this table, that information is shown in column AH.

**Table S4.** RNA-Seq: Total RNA results, two-fold change or better, $Hfq_{1-65}$/WT. This table includes all RNA species that are at least two-fold up (sheet S4A) or two-fold down (sheet S4B) for $Hfq_{1-65}$/WT total RNAs. All RNA species that are enriched more than two-fold for Hfq co-IP/total are highlighted in yellow (columns A and V). Novel sRNAs recently described in the literature or suggested by the work here, but not annotated as sRNAs in the original annotation file are indicated here as sRNA* in column B, and in the cases in which they have now been given gene names, that name is shown in column A.

**Table S5.** RNA-Seq: Hfq co-IP, two-fold change or better, $Hfq_{1-65}$/WT. This table includes all RNA species that are at least two-fold up (Sheet S5A) or two-fold down (sheet S5B) for $Hfq_{1-65}$/WT co-IP values. All RNA species that are enriched more than two-fold for Hfq co-IP/total are highlighted in yellow (columns A and V). Novel sRNAs recently described in the literature or suggested by the work here, but not annotated as sRNAs in the original annotation file are indicated here as sRNA* in column B, and in the cases in which they have now been given gene names, that name is shown in column A.

**Table S6.** Summary of literature on *in vivo* roles of *E. coli* Hfq CTD

| Reference | Year | In vivo conditions | Assays | Results | Comparison and comments | Near core/linker needed? |
|---|---|---|---|---|---|---|
| Tsui et al (8) | 1994 | *hfq1*::kmR disrupts; *hfq2*::kmR inserts after aa 78. | 1) Growth, LB, low temperature; 2) high osmolarity sensitivity | Stationary phase phenotypes (RpoS dependent) not disrupted by *hfq2*. | Original assay of Hfq roles *in vivo*. Hfq2 considered wild-type control. | Not tested here. |
| Sonnleitner et al (9) | 2004 | R66 amber, from plac in pACYC compared to WT. | 1) Qbeta growth 2) *ompA-lacZ* 3) DsrA stability | R66 stop functional for these assays. | Consistent; *ompA-lacZ* Class I regulation. DsrA consistent | Not distinguished. |
| Vecerek et al (10) | 2008 | Plac-hfq$_{1-65}$ in pACYC; reporters also plac induced and normalized to RNA levels. Levels of Hfq not compared to chromosome. | 1) Long term survival, downshift 2) Growth on succinate + DIP 3) *hfq-lacZ* autoregulation 4) *sodB-lacZ* regulation 5) RpoS Western | Hfq$_{1-65}$ fully defective in all assays. | Inconsistent; Hfq$_{1-65}$ more likely to aggregate on overproduction? | Not distinguished. |
| Olsen et al (11) | 2010 | Plac-Hfq$_{1-69}$, $_{1-72}$, $_{1-65}$, $_{1-66}$; low copy plasmid, measured as 2x chromosome. Overproduced sRNAs (pBADMicA, RybB) | 1) RpoS Western 2) RyhB sRNA, *sodB* mRNA, after DIP. 3) *ompA* mRNA with pBAD-MicA 4) *ompC* mRNA, with pBAD-RybB 5) MicM (ChiX), *ybfM* | Hfq$_{1-69}$, $_{1-72}$ generally functional. Hfq$_{1-65}$, $_{1-66}$, function for rpoS and ChiX regulation. No quantitation | Consistent; Modestly lower MicA, RybB. Modestly lower ChiX in Hfq$_{1-65}$, Hfq$_{1-66}$, not Hfq$_{1-69}$; stationary phase, consistent. Regulation normal (not quantitated). | ChiX levels dependent on near core region. |
| Beich-Frandsen et al (12) | 2011 | Hfq$_{1-65}$, $_{1-75}$, $_{1-85}$ from plac/pACYC derivatives | 1) RpoS Western, 22°C. | RpoS absent in Hfq$_{1-65}$, $_{1-75}$; present in $_{1-85}$. | Inconsistent; region between aa 75-85 implicated in | Role for near-core and other parts of CTD for RpoS. |

| | | | | | | |
|---|---|---|---|---|---|---|
| | | (pAH65, 75, 85), as per Vecerek. | | | RpoS induction.<br>If aggregate with overproduction, tip/linker helps protect. | |
| Salim et al (13) | 2012 | $Hfq_{1-65}$, $Hfq_{1-72}$, $Hfq_{1-87}$ from $P_{tac}$-Kan (pSC101 plasmid, moderate copy number) | 1) GlmS activation by GlmZ or GlmY | $Hfq_{1-72}$, $Hfq_{1-87}$ like WT; $Hfq_{1-65}$ reduced activation | Consistent; Distal face/activation defect with near-core only. | Near core needed for GlmS activation. |
| Caillet et al (14) | 2014 | $Hfq_{1-65}$, native promoter on pTX plasmid (Tsui et al) | 1) Growth<br>2) *hfq-lacZ* autoregulation<br>3) *rpoS* activation, ArcZ, DsrA levels<br>4) *oppA* repression, GcvB<br>5) *ptsG* repression, SgrS levels | Partial defect seen for auto-regulation; Other assays show no significant defect. | Consistent; Autoregulation likely dependent on strong distal binding. Modest decrease in ArcZ (processed), DsrA processed differently. | Not distinguished. |
